# Identification and characterization of two drug-like fragments that bind to the same cryptic binding pocket of *Burkholderia pseudomallei* DsbA

**DOI:** 10.1101/2021.03.25.436878

**Authors:** Guillaume A. Petit, Biswarajan Mohanty, Róisín M. McMahon, Stefan Nebl, David H. Hilko, Karyn L. Wilde, Martin J. Scanlon, Jennifer L. Martin, Maria A. Halili

## Abstract

<u>D</u>i<u>S</u>ulfide <u>B</u>ond forming proteins (DSB) play a crucial role in the pathogenicity of many Gram-negative bacteria. Disulfide bond protein A (DsbA) catalyzes the formation of disulfide bonds necessary for the activity and stability of multiple substrate proteins, including many virulence factors. Hence, DsbA is an attractive target for the development of new drugs to combat bacterial infections. Here, we identified two fragments - **1** (bromophenoxy propanamide) and **2** (4-methoxy-*N*-phenylbenzenesulfonamide), that bind to the DsbA from the pathogenic bacterium *Burkholderia pseudomallei*, the causative agent of melioidosis. Crystal structures of the oxidized *B. pseudomallei* DsbA (termed BpsDsbA) co-crystallized with **1** or **2** suggests that both fragments bind to a hydrophobic pocket that is formed by a change in the side chain orientation of tyrosine 110. This conformational change opens a “cryptic” pocket that is not evident in the *apo*-protein structure. This binding location was supported by 2D-NMR studies which identified a chemical shift perturbation of the tyrosine 110 backbone amide resonance of more than 0.05 ppm upon addition of 2 mM of fragment **1** and over 0.04 ppm upon addition of 1 mM of fragment **2**. Although binding was detected by both X-ray crystallography and NMR, the binding affinity (K_D_) for both fragments was low (above 2 mM), suggesting weak interactions with BpsDsbA. This conclusion is also supported by the modelled crystal structures which ascribe partial occupancy to the ligands in the cryptic binding pocket. Small fragments such as **1** and **2** are not expected to have high binding affinity due to their size and the relatively small surface area that can be involved in intermolecular interactions. However, their simplicity makes them ideal for functionalization and optimization. Identification of the binding sites of **1** and **2** to BpsDsbA could provide a starting point for the development of more potent novel antimicrobial compounds that target DsbA and bacterial virulence.

**Synopsis:** Describes the binding properties of two drug-like fragments to a conformationally dynamic site in the disulfide-bond forming protein A from *Burkholderia pseudomallei*.

## 1. Introduction

Fragment based drug discovery (FBDD) is the process of testing small molecules, termed fragments, against a target protein or enzyme, to identify hits that bind and - in ideal circumstances - alter the protein’s activity. Fragments often bind with low affinity due to their small size and therefore form few interactions with the protein. However, the combination and/or modification of these simple building blocks can lead to potent compounds (Murray & Rees, 2009, Woods *et al*., 2016, Kirsch *et al*., 2019). Here, we screened our fragment library against *Burkholderia pseudomallei* Disulfide bond forming protein A (BpsDsbA) using nuclear magnetic resonance (NMR) spectroscopy and X-ray crystallography. This enabled us to obtain structural information on the binding site and binding interactions between the fragment ligands and the protein.

The oxidoreductase disulfide bond forming protein A (DsbA) is required for the correct folding of multiple virulence factors, such as the type three secretion system, diverse proteases, flagellar proteins and many other virulence-associated proteins in bacteria (Heras *et al*., 2009, Coulthurst *et al*., 2008, Ireland *et al*., 2014, Bocian-Ostrzycka *et al*., 2017, Smith *et al*., 2016). A deletion of the DsbA gene (*ΔdsbA*) is not lethal for bacteria such as *Escherichia coli* (Bardwell *et al*., 1991), *Shigella flexneri* (Yu, 1998), *Francisella tularensis* (Qin *et al*., 2011, Ren *et al*., 2014) and *B. pseudomallei* (Ireland *et al*., 2014), although mutants display phenotypes such as reduced motility, reduced adhesion, and a decreased ability to replicate inside a host. Many of these phenotypes are due to the misfolding of a disulfide-containing protein in the absence of DsbA. These characteristics make DsbA an attractive target for anti-virulence drug discovery, a strategy that aims to disarm rather than kill bacteria. Such a strategy may be beneficial in reducing the selective pressure for resistance development (Allen *et al*., 2014, Heras *et al*., 2015, Mühlen & Dersch, 2016, Smith *et al*., 2016, Bocian-Ostrzycka *et al*., 2017).

*B. pseudomallei* is a Gram-negative bacterium, found predominantly in tropical areas, and the causative agent of the deadly disease melioidosis (Wiersinga *et al*., 2018). Infections by this pathogen often result in severe illness or death, even after intensive antibiotic treatment (Dance, 2014, Schweizer, 2012, Rhodes & Schweizer, 2016). *B. pseudomallei* is intrinsically resistant to many currently available antibiotics so that treatment of infection is prolonged and expensive, often requiring intravenous antibiotics for up to two weeks followed by oral antibiotics for several months (Currie, 2015).

Deletion of *dsbA* or *dsbB* results in attenuation of *B. pseudomallei* virulence, and the deletion mutants have reduced protease activity and reduced motility. Importantly, mice infected with the deletion mutants have significantly increased survival rates in infection models, compared to mice infected with wild type *B. pseudomallei* (Ireland *et al*., 2014, McMahon *et al*., 2018).

BpsDsbA is an oxidoreductase enzyme that has been biochemically characterized and its structure determined to a resolution of 1.9 Å (Ireland *et al*., 2014). The structure revealed a relatively featureless active site surface with shallow pockets and a significantly shortened hydrophobic groove (Ireland *et al*., 2014, McMahon *et al*., 2014) suggesting that it may be challenging to find small molecule inhibitors of BpsDsbA.

Techniques such as NMR, surface plasmon resonance (SPR) (Adams *et al*., 2015), and crystallography (Smith *et al*., 2016, Duncan *et al*., 2019) have all been used to identify small molecules that bind to EcDsbA, and some of these small molecules also inhibit EcDsbA in activity based assays (Halili *et al*., 2015, Totsika *et al*., 2018, Mohanty *et al*., 2017). Although inhibitors and small molecules screening has mostly focused on EcDsbA, there has been some success in identifying molecules binding to BpsDsbA as well (Nebl *et al*., 2020, McMahon *et al*., 2018). A short peptide derived from the sequence of its partner protein BpsDsbB has been shown by crystallography to bind BpsDsbA, revealing a relatively flat interaction site around the active site of the protein (McMahon *et al*., 2018). Additionally a fragment was shown to bind at a conformationally dynamic site on the surface of the protein, using NMR (Nebl *et al*., 2020).

In this work we report two fragments that bind to BpsDsbA, which could potentially be suitable for further development as inhibitors. These are bromophenoxy propanamide (**1**), and 4-methoxy-*N*-phenylbenzenesulfonamide (**2**). Binding was characterized using NMR and X-ray crystallography. Both **1** and **2** bind to a cryptic pocket on BpsDsbA - not observed in the *apo*-BpsDsbA structure - located adjacent to the redox active site. This transient (or “cryptic”) pocket is formed by a shift in the side chain conformation of a tyrosine residue to accommodate the fragments.

## 2. Materials and methods

### 2.1 Protein expression and purification for crystallization and peptide oxidation assay

Recombinant BpsDsbA was expressed as described in Ireland *et al*. (Ireland *et al*., 2014). Briefly, plasmids with the BpsDsbA gene in a modified pET22 vector with Tobacco etch virus protease (TEV) cleavage site followed by a His_6_ metal affinity tag were transformed into *E. coli* B21(DE3)pLysS competent cells, grown in 10 ml Lysogeny broth (LB) containing chloramphenicol (CAM) and ampicillin (AMP) and incubated at 37°C overnight. Pre-cultures were used to start 1 l culture in autoinduction media also containing CAM and AMP (Studier, 2005). pET28a plasmid containing the BpsDsbB gene with a non-cleavable His_8_-tag was used to transform C41 *E. coli* cells specialized in membrane protein expression, also using autoinduction media supplemented with kanamycin (KAN).

BpsDsbA was purified according to the protocol described in Ireland *et al*. (Ireland *et al*., 2014). In short, after expression cells were harvested by centrifugation at 6000 x *g*. The pellet was resuspended in buffer containing 25 mM tris(hydroxymethyl)aminomethane (TRIS) pH 7.5 and 150 mM NaCl. Cells were lysed by two passages at 165 MPa in a cell disrupter (Constant System) and the debris separated from supernatant containing the soluble protein by centrifugation (30 min at 30,000 x *g*). Imidazole (pH 7.5) was then added to the supernatant to a final concentration of 5 mM and the solution was subjected to immobilized metal affinity chromatography (IMAC) by incubation with TALON cobalt resin (Takara) for 1 hr at 4°C. Resin-bound protein was loaded onto a gravity flow column and washed with 2 x 5 column volumes (CV) of wash buffer (10 mM imidazole, 500 mM NaCl and 25 mM TRIS pH 7.5) before elution in 5 CV of 300 mM imidazole, 150 mM NaCl and 25 mM TRIS pH 7.5. The protein was buffer exchanged to remove imidazole using a 16/260 HiLoad desalting column (GE healthcare). BpsDsbA was then incubated with TEV protease in a 1:50 (TEV:BpsDsbA) stoichiometric ratio overnight at 4°C. The next day the cleaved His_6_-tags, non-cleaved protein and the TEV protease (also His_6_-tagged) were removed by reverse IMAC in TALON resin with the target protein in the flowthrough. The protein was oxidized by mixing with a molar excess of oxidized glutathione (GSSG) at room temperature for 1 hr (50:1 stochiometric ratio of GSSG:BpsDsbA), the oxidation state of the protein was monitored by Ellman test (Ellman, 1959). A final size exclusion step in 25 mM (4-(2-hydroxyethyl)-1-piperazineethanesulfonic acid (HEPES) pH 7.5 and 150 mM NaCl was used to remove GSSG and impurities. The fractions corresponding to the protein were pooled, concentrated to 33 mg/ml using an Amicon Ultra 50 ml 10 kDa molecular weight cut-off centrifugal filter (Merck Millipore) and then aliquoted before flash freezing the protein sample in liquid nitrogen. Protein concentration was estimated using a NanoDrop™ ND-1000 (Thermo Scientific) photo-spectrometer.

Membrane preparations of BpsDsbB for the peptide oxidation assay were generated using a method similar to that reported in (Christensen *et al*., 2019). Briefly, the gene for BpsDsbB (Uniprot ID Q63RY4) was inserted in a pET28a plasmid in front of a sequence coding for a C-terminal non-cleavable His_8_-tag. The plasmid was inserted in C41 *E. coli* cells specialized for the expression of membrane proteins (Wagner *et al*., 2008) which were grown in autoinduction medium (Studier, 2005) for 24 hr at 30 °C with shaking at 220 rpm. The cells were harvested by centrifugation at 6,000 x *g* and resuspended in phosphate buffered saline (PBS). Cells were disrupted by two passages at 207 MPa through a cell disruptor (Constant System). Large debris was removed by centrifugation for 15 min at 15,000 x *g*, and membranes containing protein were further separated from solution by ultracentrifugation for 1 hr 15 min at 180,000 x *g*. The membrane pellet was resuspended in PBS prior to usage in peptide oxidation assay.

### 2.2 Expression and purification of [U-^15^N]-labelled BpsDsbA for NMR spectroscopy

Uniformly ^15^N-labelled ([U-^15^N]) BpsDsbA was expressed at the National Deuteration Facility (NDF), Australian Nuclear Science and Technology Organization (ANSTO). The gene encoding BpsDsbA was inserted in a pET24a vector maintaining the TEV protease cleavable N-terminal His_6_-tag for protein expression using a high cell density protocol as reported previously (Duff *et al*., 2015). Briefly, 300 μl of freshly transformed *E. coli* BL21Star™(DE3) cells were inoculated into 10 ml of H_2_O ModC1 minimal medium and incubated overnight at 30 °C with shaking at 220 rpm. This cell suspension was diluted 5x in fresh ^1^H, ^15^N-ModC1 medium (40 g/l glycerol, 5.16 g/l ^15^NH_4_Cl ≥ 98 atom % ^15^N) and grown at 37 °C for two OD_600_ doublings. Finally, cells were inoculated into fresh ^1^H, ^15^N-ModC1 to a volume of 100 ml and grown to an OD_600_ of 0.9 before inoculation into 900 ml of labelled expression medium as described in a 1 l working volume bioreactor. *E. coli* cells were grown at 25 °C until OD_600_ reached 14.8 and expression induced by addition of isopropylthio-β-D-galactopyranoside (IPTG) at a final concentration of 1 mM. After 22.5 hr induction at 20 °C during which a further 5.1 g of ^15^NH_4_Cl was added to the culture, the labelled cell suspension was pelleted by centrifugation at 8000 x *g* for 20 min and the pellet stored at −80 °C.

BpsDsbA purification was performed in-house using the protocol reported previously by Nebl *et. al*., (Nebl *et al*., 2020). Briefly, the frozen cell pellet was resuspended in lysis buffer comprising 50% BugBuster MasterMix (Novagen) and 50% buffer A containing 20 mM HEPES, 100 mM NaCl, 10 mM imidazole (pH 8.0) at 2.5 ml/g of cell pellet. One EDTA-free protease inhibitor tablet (Roche) was added into the lysis buffer to prevent proteolysis. The mixture was agitated for 30 min at room temperature. To ensure complete cell lysis, sonication was performed on ice for 30 s x 7 times at 50% duty cycle. The lysate was centrifuged at 75,465 x *g* for 30 min at 4°C. The supernatant was filtered through a 0.22 μm syringe filter and loaded onto an immobilized Ni^2+^ affinity column (HisTrap HP 5ml, GE Healthcare) using buffer A and eluted using a gradient of 10-500 mM imidazole. Fractions containing target protein were pooled and exchanged back to 100% buffer A using a Sephadex desalting column (HiPrep 26/10 column, GE Healthcare). TEV cleavage was performed overnight at 23°C with 1 mM DTT, and 0.1 mg TEV per 10 mg protein. A second reverse IMAC step was performed to collect the TEV-cleaved protein and remove His-tagged TEV protease, cleaved His_6_-tag and uncleaved BpsDsbA. The TEV-cleaved BpsDsbA was oxidized overnight at 4 °C using freshly prepared copper-phenanthroline at a final concentration of 1.5 mM. A final desalting step was performed to remove copper-phenanthroline and exchange the sample into 50 mM HEPES, 50 mM NaCl, 2 mM EDTA (pH 6.8) prior to purification by size exclusion chromatography using a gel filtration column (HiLoad 26/60 Superdex 75 column, GE Healthcare). The sample was concentrated using a 10 kDa molecular weight cut-off centrifugal filter (Merck Millipore). The protein concentration was estimated using a NanoDrop™ 1000 (Thermo Scientific) spectrophotometer. Finally, 1 mM phenylmethylsulfonyl fluoride (PMSF), 0.02 % NaN3 and 10% D_2_O were added into the protein stock prior to NMR experiments.

### 2.3 Acquisition of small molecule fragments

Fragment bromophenoxy propenamide (**1**) (≥95% purity) was purchased from hit2lead (Chembridge Corporation, San Diego, CA).

Fragment 4-methoxy-*N*-phenylbenzenesulfonamide (**2**) was synthesized according to literature procedure (Bernar *et al*., 2018). See supporting information for more details (Figure S1).

### 2.4 Quality control and solubility assessment of 1 and 2 in aqueous NMR buffer

The solubility of **1** and **2** was assessed by recording a set of 1D ^1^H-NMR spectra in aqueous NMR buffer (50 mM HEPES, 25 mM NaCl, 2 mM EDTA, 2% D_6_-DMSO, 100 μM DSS, 10% D_2_O at pH 6.8). Chemical shift and peak volumes of individual proton signals in the 1D ^1^H spectra were measured in order to identify possible aggregation either via concentrationdependent changes in the chemical shifts of the peaks or deviation from the expected concentration-dependent increase in peak volume (LaPlante *et al*., 2013). 1D ^1^H spectra were collected on a 600 MHz spectrometer equipped with CryoProbe at 298 K with a relaxation delay of 10 s. 1D ^1^H spectra were processed and analyzed by Mnova (Bernstein *et al*., 2013).

### 2.5 Chemical shift perturbation analysis and estimation of ligand binding affinity (K_D_) by 2D [^15^N,^1^H]-HSQC NMR

Binding affinity of **1** and **2** against oxidized BpsDsbA was assessed by titration against 100 μM ^15^N-labelled BpsDsbA. Backbone assignments of both redox states of BpsDsbA have been reported previously by Nebl *et. al*., (Nebl *et al*., 2020), these assignments were used for the chemical shift perturbations (CSP) analysis in the 2D [^15^N,^1^H]-heteronuclear single quantum coherence (HSQC) spectra using either CARA ((Keller, 2005), Diss. ETH Nr. 15947; http://cara.nmr.ch/) or SPARKY (Lee *et al*., 2015). Fragments **1** and **2** were titrated at concentrations of 0.25, 0.50, 1 and 2 mM with 100 μM [U-^15^N]-BpsDsbA in NMR buffer (50 mM HEPES, 25 mM NaCl, 2 mM EDTA, 2% D_6_-DMSO, 1 mM PMSF, 10% D_2_O at pH 6.8). CSPs were calculated for each perturbed peak according to equation 1 (Nebl *et al*., 2020).

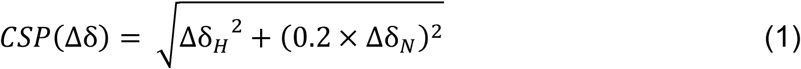

where Δδ_H_ and Δδ_N_ are the measured differences between the chemical shifts in the free vs bound spectra for the hydrogen and nitrogen signal (in ppm), respectively. In an effort to estimate the dissociation constants (*K*_D_) of fragment **1** and fragment **2** the CSP titration data were fitted to a one-site binding model in GraphPad Prism using nonlinear regression with equation 2 (Nebl *et al*., 2020).

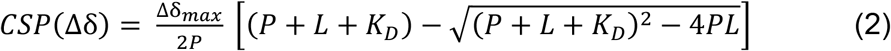

where P and L are total concentrations of protein and ligand respectively, Δδ_max_ is the maximum CSP upon saturation and *K*_D_ is the calculated dissociation constant. However, the CSP responses were observed to increase linearly with concentration and reliable estimates of *K*_D_ could not be obtained. These data do provide an indication of the site of interaction between the ligand and oxidized BpsDsbA, by plotting the CSP magnitude as a gradient onto the crystal structure of BpsDsbA.

### 2.6 Crystallization of BpsDsbA for soaking experiments

Oxidized BpsDsbA, purified in 25 mM HEPES with 150 mM NaCl was concentrated to 25-33 mg/ml and dispensed in 100 nl protein onto a MRC-2 sitting drop 96 well plate (Hampton research) and mixed with 100 nl of crystallization buffer (0.1 M HEPES pH 7.5, 0.2 M Li_2_SO_4_ and gradient of PEG3350 28-34%). Crystal needles typically appeared after several hours and continued to grow for 4-5 days. Fragments were dissolved in DMSO to a final concentration between 5 mM and 25 mM. The fragment-DMSO solution was mixed with the crystallization buffer to final concentrations ranging from 0.25 mM to 1.25 mM and a BpsDsbA crystal was soaked in the fragment solution for approximatively 2 hr. Similarly, crystals used to generate the background <u>Pan</u>-<u>d</u>ataset <u>d</u>ensity <u>a</u>nalysis (PanDDA) (Pearce *et al*., 2017) map were soaked in mother liquor containing 5% DMSO without fragment for 2 hrs. After soaking, crystals were fished using nylon loops and cryo-cooled in liquid nitrogen (the high concentration of PEG in the mother liquor acted as a cryoprotectant).

### 2.7 Co-crystallization of BpsDsbA with 1 or with 2

BpsDsbA was purified and oxidized as described above, concentrated to 33 mg/ml and mixed with 10 mM of **1** and kept on ice for 2 hr. The solution was centrifuged to remove excess fragment that did not dissolve. A 100 nl drop of solution containing the protein in the presence of **1** was then dispensed in hanging drop and combined with a 100 nl drop of mother solution from commercial screens at 20°C by a Mosquito robot (SPT-Labtech). Crystal needles grew in 60% tacsimate after a few hours and kept growing over 2-3 days. A large needle crystal was fished with a nylon loop and cryoprotected in mother liquor containing 20% ethylene glycol. The crystal was then cryo-cooled in liquid nitrogen and tested in X-ray diffraction experiment. Similarly, oxidized BpsDsbA at 33 mg/ml was mixed with a large molar excess of **2** and incubated on ice for 2 hrs. Once again the solution was centrifuged to remove excess fragment that did not dissolve. A 100 nl drop of solution containing protein in the presence of **2**, was mixed with 100 nl of crystallization solution, dispensed as a hanging drop onto MRC-2 crystallization plate and incubated at 20°C. Long crystal needles appeared after 2-3 days in 0.1 M HEPES pH 7.5, 0.2 M Li_2_SO_4_ and 29.5% PEG3350. Crystals were fished with nylon loops and flash frozen without additional cryoprotection.

### 2.8 X-ray diffraction experiments and refinement

X-ray diffraction data were collected at the Australian Synchrotron (part of the Australian Nuclear Science and Technology Organization (ANSTO)) on the macromolecular crystallography beamlines MX1 (ADSC Quantum 210r Detector) and MX2 (Eiger 16M detector, funded by the Australian Cancer Research Foundation). Data were indexed, scaled and analyzed with the autoProc pipeline (Vonrhein *et al*., 2011) where possible - or manually with XDS (Kabsch, 2010) when autoProc analysis failed. Structures were solved by molecular replacement using the oxidized BpsDsbA model PDB ID 4K2D and refined using the Dimple pipeline, part of CCP4 (Winn *et al*., 2011). Occasionally datasets required a step wise analysis, in which case phaser (McCoy *et al*., 2007) and phenix.refine (Adams *et al*., 2010) were used. Structures were then manually inspected with Coot (Emsley *et al*., 2010) and Molprobity (Chen *et al*., 2010). Refinement steps were repeated as required, alternating between Coot and phenix.refine. Ligand coordinates were generated from SMILES files using eLBOW (Moriarty *et al*., 2009). Initial inspection of the datasets did not suggest density indicative of ligand binding. PanDDA (pandda.analyse) was run on the Griffith University high performance cluster (HPC) Gowonda, following the instructions from https://pandda.bitbucket.io/tutorials.html. “Hits” were inspected with PanDDA (pandda.inspect) through the coot interface. The majority of these hits were false positives. Fragments **1** and **2** were identified as hits and further refined with phenix.refine and Coot.

The structures of BpsDsbA co-crystallized with fragments **1** or **2** were solved by molecular replacement with Dimple (Winn *et al*., 2011) and phaser (McCoy *et al*., 2007) using the oxidized BpsDsbA structure (PDB ID 4K2D) as a search model. Models were refined using phenix.refine (Afonine *et al*., 2012) and Coot, and Molprobity (Chen *et al*., 2010) was used for validation.

### 2.9 Peptide oxidation assay

The ability of fragments **1** and **2** to inhibit BpsDsbA was tested in a peptide oxidation assay described previously in Halili *et al*. 2015 (Halili *et al*., 2015). Briefly, a synthetic peptide with two fluorescent groups at each extremity and two cysteines near each end can be oxidized in the presence of active DsbA. Upon oxidation the two fluorescent groups are brought into close contact and can be excited at 340 nm to fluoresce at 615 nm. During the typical uninhibited reaction, the fluorescence of the peptide increases over 10 to 15 min until a plateau is reached. In the presence of BpsDsbA inhibitors, the enzyme fails to oxidize the peptide and the fluorescence does not increase over time.

Samples were prepared in 384 well plates with a final concentration of reactants of 60 nM BpsDsbA, 1.6 μM of BpsDsbB in membranes, a range from 0 to 20 mM of fragment, and 10 μM of substrate peptide, in a final volume of 50 μl. The reaction was monitored using a Synergy H1 Hybrid plate reader (Biotek™), with the excitation wavelength set to 340 nm, emission to 620 nm and a 100 μs delay between the excitation and reading. Plates were monitored for 3 hrs, until a reaction plateau was reached.

## 3. Results

### 3.1 Identification of fragments binding to oxidized BpsDsbA using crystal soaking experiments and PanDDA analysis

An initial screen of ~1130 fragments obtained from the Monash Institute for Pharmaceutical Science (MIPS) fragment libraries (Doak *et al*., 2014) was performed using ligand-detected saturation transfer difference (STD) NMR (Mayer & Meyer, 1999) against the oxidized BpsDsbA and EcDsbA proteins (Nebl *et al*., 2020, Adams *et al*., 2015). A set of fragments was initially identified as binding to BpsDsbA by STD-NMR. These hits were considered validated if they elicited detectable CSP in protein detected 2D [^15^N,^1^H]-HSQC spectra of BpsDsbA (Nebl *et al*., 2020). Among these promising candidates, a small subset of fragments was selected for further analysis in this study.

A total of 29 unique fragments (Figure S2) were dissolved separately in 100% DMSO at a concentration up to 25 mM and the solution was used to soak individual BpsDsbA crystals. Crystals were exposed to X-rays either at the Australian Synchrotron (on the MX1 or MX2 beamlines) or on the laboratory source at The University of Queensland UQROCX crystallization facility. All the crystals diffracted in the P2_1_2_1_2_1_ space group, all unit cell angles were 90° as expected for this space group. All the axes length were found between 59.0 and 60.0 Å for axis A, 61.5 to 63.5 Å for axis B and 68.0 to 70.5 Å for axis C. There was no interpretable positive difference Fourier density in any of the datasets to indicate binding of the different fragments to the protein. We then reprocessed the diffraction datasets using a more sensitive method, PanDDA (Pearce *et al*., 2017). We generated a background map from 32 X-ray diffraction datasets of the *apo*-protein soaked in DMSO (resolution ranging from 1.70 Å to 2.28 Å). This was used as the “ground-state” model to reanalyze datasets of the protein soaked with the individual fragments. Using this method, we identified that two of the soaked crystal datasets showed peaks of positive Fourier densities that could be interpreted as bound fragments **1** and **2** (structure of fragments shown in Figure S2 and S3), respectively. In both models, the fragments bind near tyrosine 110 (Y110), causing a change in the tyrosine sidechain position in comparison to the *apo*-structure (Figure 1B). This shift revealed the presence of a small hydrophobic pocket into which each fragment binds (Figure 1C and D). The binding of both fragment to BpsDsbA was then reproduced using independent co-crystallization experiments.

**Figure 1 –.**
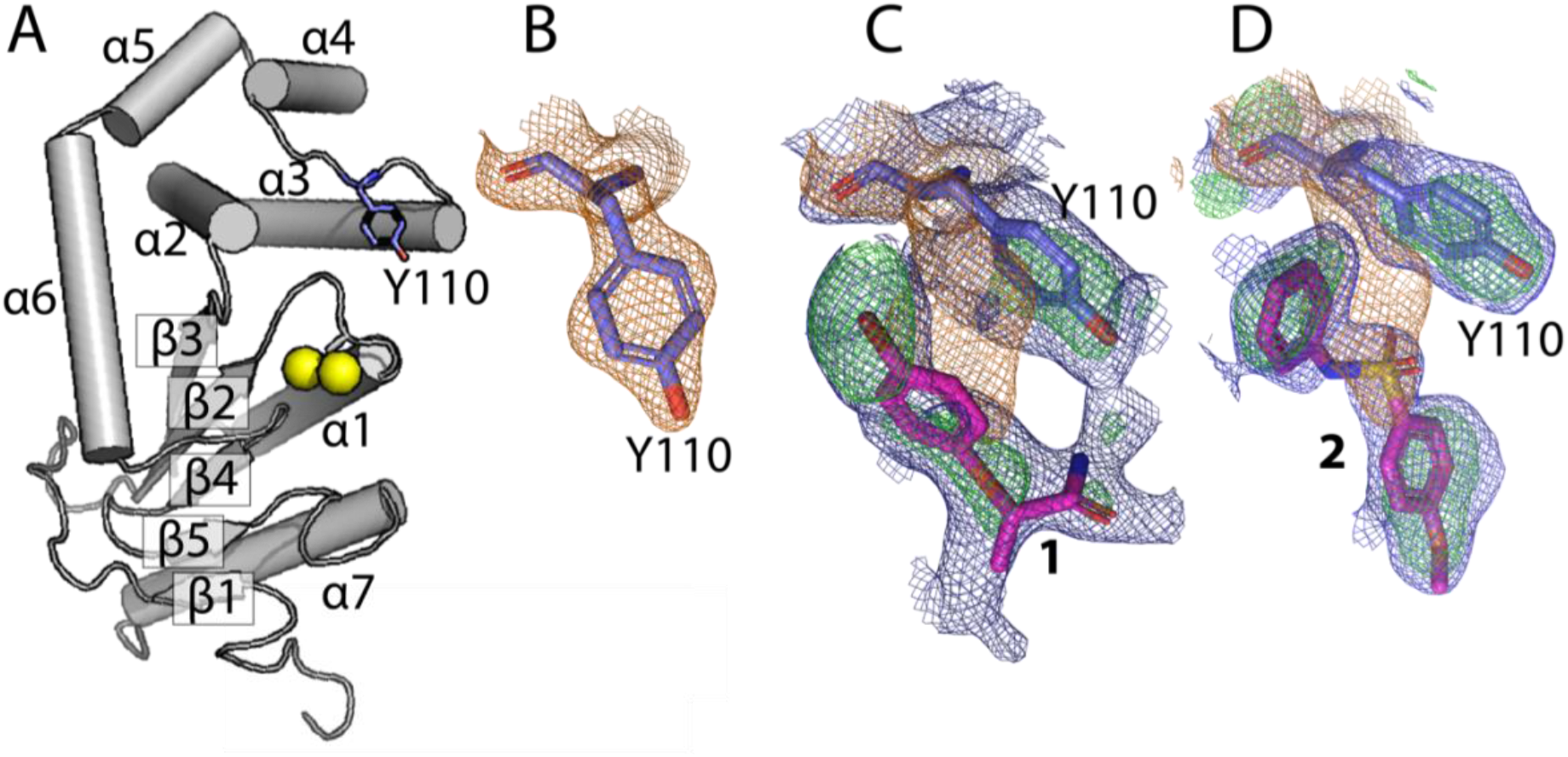
Event map generated by PanDDA around Y110 and fragments 1 and 2. **(A)** Architecture of the *apo*-BpsDsbA structure (PDB ID 4K2D, (Ireland *et al*., 2014)) represented as a cartoon. α-helices and β-strands are numbered α1-7, β1-5 respectively. The active site cysteines are indicated by yellow spheres, Y110 is represented in purple in stick format. **(B)** Close up of the orientation of Y110 in the *apo*-structure (no ligand present) and **(C)** and **(D)** in the presence of **1** and **2** respectively. In this orientation, the Y110 sidechain rotates to the right (viewed *along* the C^β^-C^γ^ bond) towards helix α3 compared to the *apo-*structure. This shift opens a small hydrophobic pocket into which each respective fragment binds. The reference *apo*-map 2Fo-Fc is contoured at 1σ and shown in orange, and is the result of averaging 32 electron density maps of *apo*-BpsDsbA. Blue is the event difference map contoured at 3σ for **1** and **2** (C and D, respectively). It displays the differences between the reference map and the map of the ligand-soaked datasets. In green, the PanDDA Z-maps highlight significant deviations between the reference map and the ligand soaked-dataset (Pearce *et al*., 2017)

### 3.2 2D [^15^N,^1^H]-HSQC-NMR binding assay of 1 and 2 with BpsDsbA

Fragments **1** and **2** were previously identified as binding to oxidized BpsDsbA in a HSQC-NMR binding assay (Nebl *et al*., 2020). The original HSQC screen was conducted using mixtures of two fragments. To confirm binding we followed up the original experiment by recording 2D [^15^N,^1^H]-HSQC of BpsDsbA with each fragment individually.

Prior to HSQC-screening of the two fragments, we evaluated their solubility in the NMR buffer (50 mM HEPES, 25 mM NaCl, 2 mM EDTA, 2% D_6_-DMSO, 100 μM DSS, 10% D_2_O at pH 6.8). This confirmed that **1** and **2** were soluble in the NMR buffer (Figures S3 and S4). Overlays of the 2D [^15^N, ^1^H]-HSQC spectra of oxidized BpsDsbA (100 μM) in the absence and presence of **1** (2 mM) and **2** (1 mM) are shown in Figures 2 and 3, respectively. Chemical shift perturbations (CSPs) resulting from the addition of **1** and **2** are mapped onto the crystal structure of oxidized BpsDsbA in Figures 2 and 3 to provide a visual estimate of their binding sites. Both fragments produced backbone amide CSPs > 0.02 ppm for residues C43, E48, H105, Y110 and L111. Two additional residues: A72 and K108, showed CSP > 0.02 ppm for **1**. These residues form a cluster between the C^43^PHC^46^ active site, the *cis*Pro loop adjacent to the active site, the C-terminal residues of helix α3, a loop connecting helix α3 and α4 and a loop between β3 and α2, connecting the two domains of the protein (Figures 4A and 4B). The location of the largest CSPs suggest that **1** and **2** may interact near the catalytic site of oxidized BpsDsbA; this site has been previously identified as a small molecule binding site (Nebl *et al*., 2020). Linear chemical shift trajectories upon increasing fragment concentrations (Figure S5) indicate that the fragments are in fast exchange on the chemical shift time scale, suggesting weak binding (Ziarek *et al*., 2011).

**Figure 2 –.**
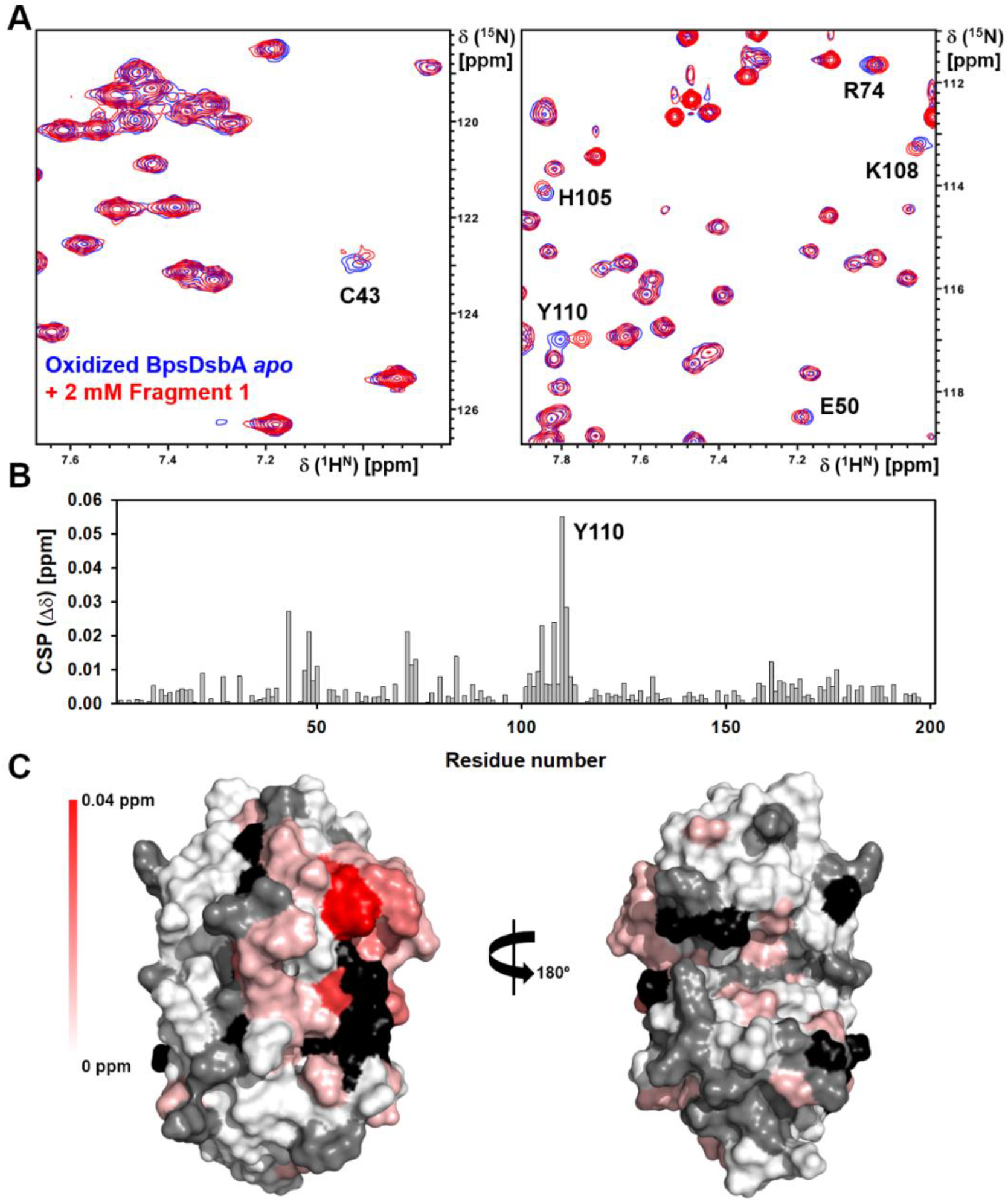
Characterization of bromophenoxy propanamide (1) binding to oxidized BpsDsbA by 2D [^15^N, ^1^H]-HSQC NMR. (**A**) Expanded regions of the 2D [^15^N,^1^H]-HSQC data highlighting the backbone amide CSP for selected residues of BpsDsbA without (blue) and with 2 mM of fragment **1** (red). (**B**) CSP observed for each BpsDsbA residue. (**C**) CSPs resulting from the addition of 2 mM fragment **1** are mapped onto the crystal structure of oxidized BpsDsbA (PDB ID 4K2D) as a color gradient from red (CSP = 0.04 ppm) to white (CSP = 0 ppm). Non-shifting residues are shown in grey. Residues with unassigned amides and proline residues are shown in black. N-terminal residues (A1-G14) were removed for clarity.

**Figure 3 –.**
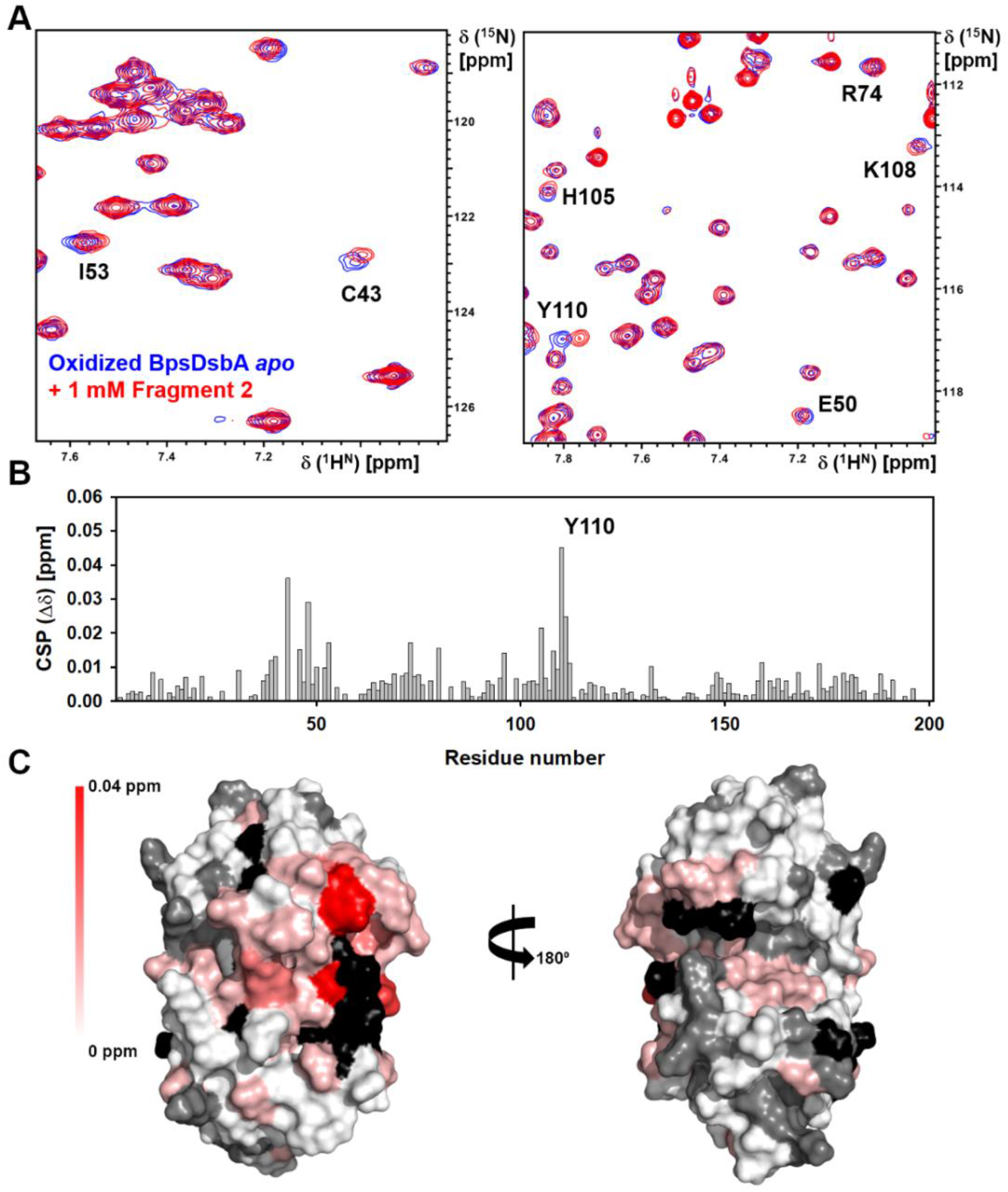
Characterization of methoxy-phenylbenzenesulfonamide (2) binding to oxidized BpsDsbA by 2D [^15^N, ^1^H]-HSQC NMR. **(A)** Expanded regions of the 2D [^15^N,^1^H]-HSQC data highlighting the backbone amide CSP for selected residues of BpsDsbA without (blue) and with 1 mM of fragment **2** (red). **(B)** CSP observed for each BpsDsbA residue. **(C)** CSPs resulting from the addition of 1 mM **2** are mapped onto the crystal structure of oxidized BpsDsbA (PDB ID 4K2D) as a color gradient from red (CSP = 0.04 ppm) to white (CSP = 0 ppm). Residues with unassigned amides and proline residues are shown in black. Non-shifting residues are shown in grey. N-terminal residues (A1-G14) were removed for clarity.

**Figure 4 –.**
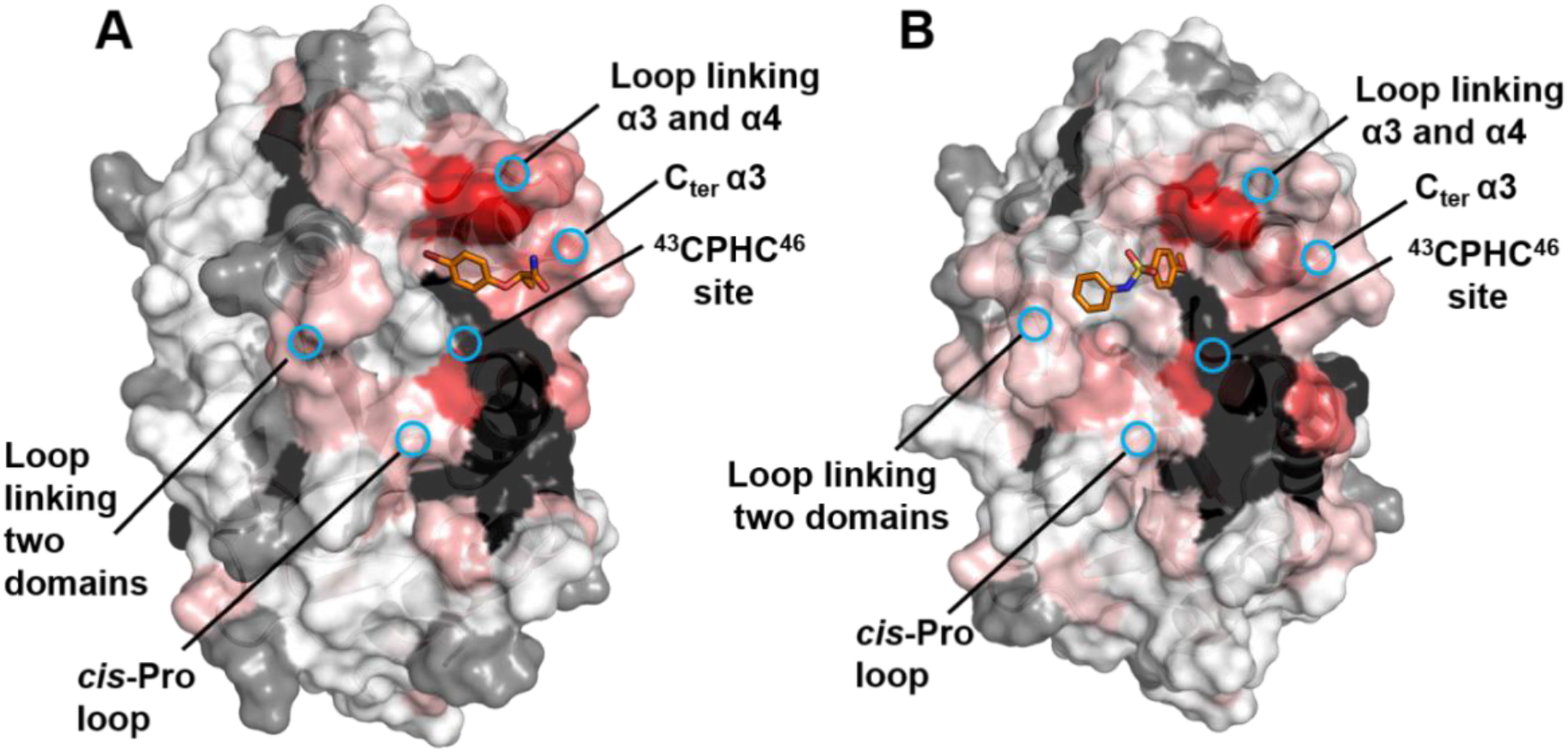
Characterization of fragment 1 and 2 binding to oxidized BpsDsbA by 2D [^15^N, ^1^H]-HSQC NMR. Chemical shift perturbations (CSP) resulting from addition of 2 mM **1** (**A**) and 1 mM **2** (**B**) are mapped onto their corresponding complex crystal structures (PDB ID 7LUH for **1** and chain A of PDB ID 7LUJ for **2**). CSPs are plotted as a color gradient from red (CSP = 0.04 ppm) to white (CSP = 0 ppm). Residues with unassigned amides and proline residues are shown in black. Non-shifting residues are shown in grey. N-terminal residues (A1-G14) were removed for clarity.

To estimate the binding affinity of fragments **1** and **2** with oxidized BpsDsbA, we recorded a series of [^15^N,^1^H]-HSQC spectra of 100 μM BpsDsbA with increasing concentrations of fragment **1** (0 – 2 mM) and **2** (0 – 1 mM). For both fragments, CSPs were observed to increase linearly with respect to concentration, and saturation was not achieved. Figure S6 shows the concentration-dependent CSP profiles of several binding site residues. The CSP did not reach saturation at 2 mM ligand concentration, indicating that fragments **1** and **2** bind weakly with a K_D_ greater than the highest concentrations tested.

We previously observed redox dependent ligand binding to BpsDsbA, and we hypothesized that this is due to differences in the dynamics of reduced and oxidized BpsDsbA (Nebl *et al*., 2020). Here, we repeated the HSQC-titrations of **1** and **2** against reduced BpsDsbA, and we did not observe any significant CSP (Figure S7). This indicates that fragments **1** and **2** bind preferentially to the oxidized form of BpsDsbA.

### 3.3 BpsDsbA co-crystallized with bromophenoxy propanamide (1) in a cryptic pocket binding site

Oxidized BpsDsbA was co-crystallized with **1** in 60% tacsimate and the resulting crystals diffracted to a resolution of 1.84 Å on beamline MX2 at the Australian Synchrotron, in space group, P2_1_2_1_2_1_. The structure was solved by molecular replacement using the original oxidized BpsDsbA structure (PDB ID 4K2D) (Ireland *et al*., 2014) as a search model. The structure was further refined by addition of the ligand giving final R_work_ and R_free_ values of 16.82% and 19.84% respectively (Table 1), suggesting that the model is a good fit to the data. Overall, the backbone structure (Cα) of BpsDsbA in complex with **1** was very similar to that of the structure with no ligand (Figure 5), with a root mean square deviation (RMSD) of 0.14 Å between the residues of the two proteins (191 residues aligned with 191 residues, with the Pymol super function (Schödinger, 2015))

**Figure 5 –.**
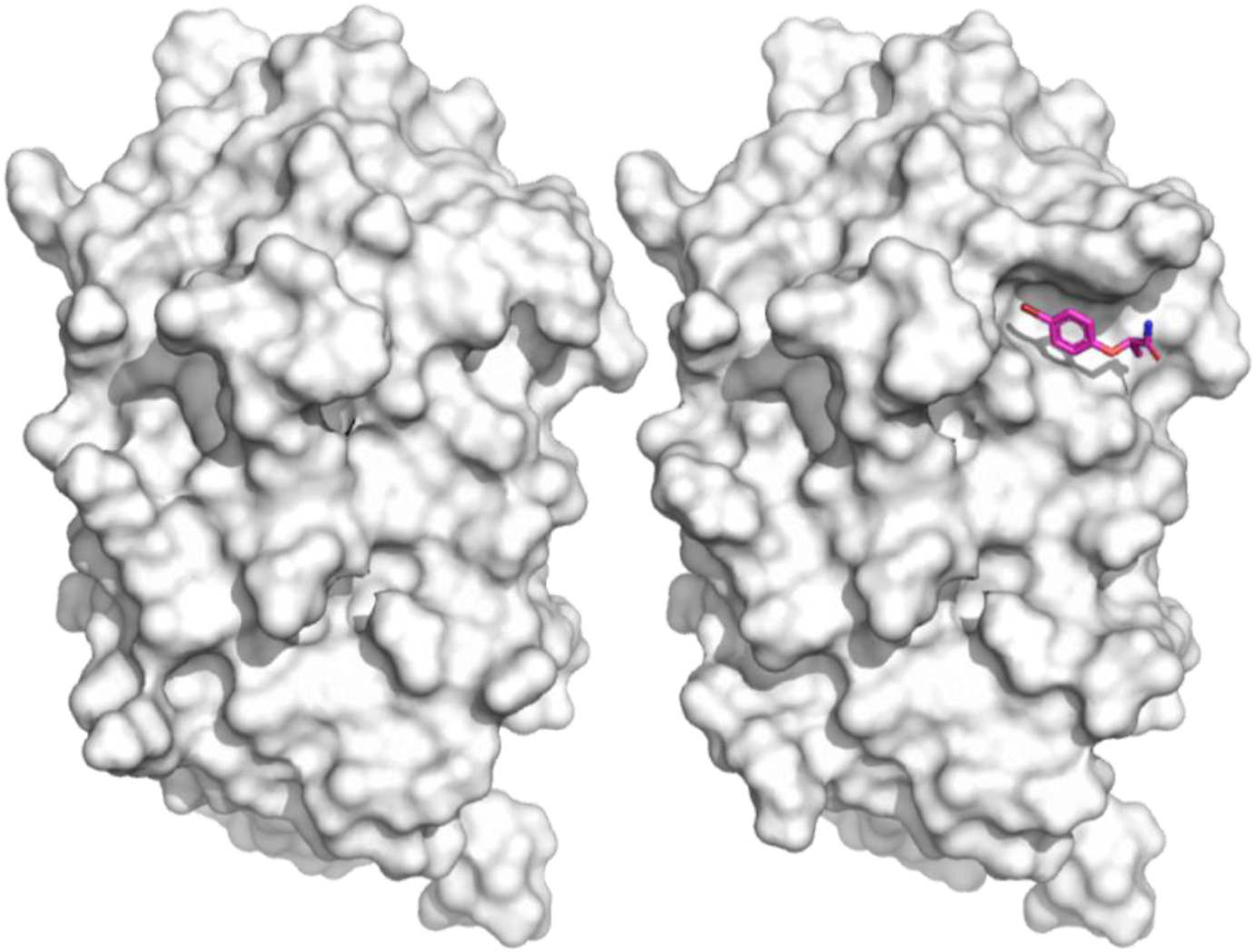
Structure of BpsDsbA co-crystallized with bromophenoxy propanamide (1). On the left, the structure of the oxidized *apo*-protein (PDB ID 4K2D). On the right, the structure of oxidized BpsDsbA in presence of **1** (PDB ID 7LUH). The proteins are shown as a white surface, with the fragment **1** shown in magenta and binding to a small pocket near Y110.

**Table 1:** Data collection and refinement statistics

|  | BpsDsbA + **1** (PDB ID 7LUH) | BpsDsbA +**2** (PDB ID 7LUJ) |
| --- | --- | --- |
| **Wavelength (Å)** | 0.9537 | 0.9537 |
| **Resolution range (Å)** | 36.7 - 1.84 (1.91 - 1.84) | 38.4 - 2.31 (2.40 - 2.31) |
| **Space group** | P 2<sub>1</sub> 2<sub>1</sub> 2<sub>1</sub> | P 2<sub>1</sub> |
| **Unit cell (Å, °)** | 59.5, 62.9, 69.4, 90, 90, 90 | 69.4 59.4 105.4 90 104.9 90 |
| **Total reflections** | 156535 (15993) | 71917 (7044) |
| **Unique reflections** | 23108 (2249) | 36201 (3064) |
| **Multiplicity** | 6.8 (7.1) | 2.0 (2.0) |
| **Completeness (%)** | 99.22 (98.38) | 97.40 (84.08) |
| **Mean I/sigma(I)** | 10.78 (0.69) | 5.20 (0.84) |
| **Wilson B-factor** | 31 | 45 |
| **R<sub>merge</sub>** | 0.02698 (0.4728) | 0.06982 (0.5694) |
| **R<sub>meas</sub>** | 0.03816 (0.6687) | 0.09874 (0.8052) |
| **R<sub>pim</sub>** | 0.02698 (0.4728) | 0.06982 (0.5694) |
| **CC<sub>1/2</sub>** | 0.999 (0.688) | 0.992 (0.508) |
| **CC\*** | 1 (0.903) | 0.998 (0.821) |
| **Reflections used in refinement** | 23062 (2248) | 35652 (3064) |
| **Reflections used for R<sub>free</sub>** | 1986 (183) | 1785 (137) |
| **R<sub>work</sub>** | 0.1682 (0.2927) | 0.2127 (0.2984) |
| **R<sub>free</sub>** | 0.1984 (0.3388) | 0.2704 (0.3234) |
| **CC<sub>work</sub>** | 0.969 (0.839) | 0.947 (0.624) |
| **CC<sub>free</sub>** | 0.955 (0.763) | 0.920 (0.561) |
| **Number of non-hydrogen atoms** | 1735 | 6108 |
| **macromolecules** | 1534 | 5838 |
| **ligands** | 13 | 72 |
| **solvent** | 188 | 224 |
| **Protein residues** | 192 | 756 |
| **RMS(bonds, Å)** | 0.008 | 0.003 |
| **RMS(angles, °)** | 0.82 | 0.53 |
| **Ramachandran favored (%)** | 98.41 | 98.79 |
| **Ramachandran allowed (%)** | 1.59 | 1.21 |
| **Ramachandran outliers (%)** | 0.00 | 0.00 |
| **Rotamer outliers (%)** | 0.60 | 0.49 |
| **Clashscore** | 1.62 | 2.77 |
| **Average B-factor (Å<sup>2</sup>)** | 41 | 53 |
| **macromolecules** | 40 | 53 |
| **ligands** | 68 | 72 |
| **solvent** | 45 | 46 |
| **Number of TLS groups** | 10 | 12 |

Statistics for the highest-resolution shell are shown in parentheses. $CC^* = \sqrt{\frac{2CC_{1/2}}{1+CC_{1/2}}}$ (Karplus & Diederichs, 2012)

The data collected for BpsDsbA + fragment **1** (PDB ID 7LUH) showed difference density corresponding to the ligand without the use of PanDDA and this was verified by using a Polder map (an OMIT map that accounts for solvent (Liebschner *et al*., 2017)). The Polder map showed positive difference density for the ligand at 3σ contour level (Figure 6). Additional unexplained density was present near the modelled carboxamide of the ligand, possibly from water or a component of the crystallization conditions (malonate, citrate, succinate, malic acid, acetate, formate and tartrate). We chose not to model anything into this density.

**Figure 6 –.**
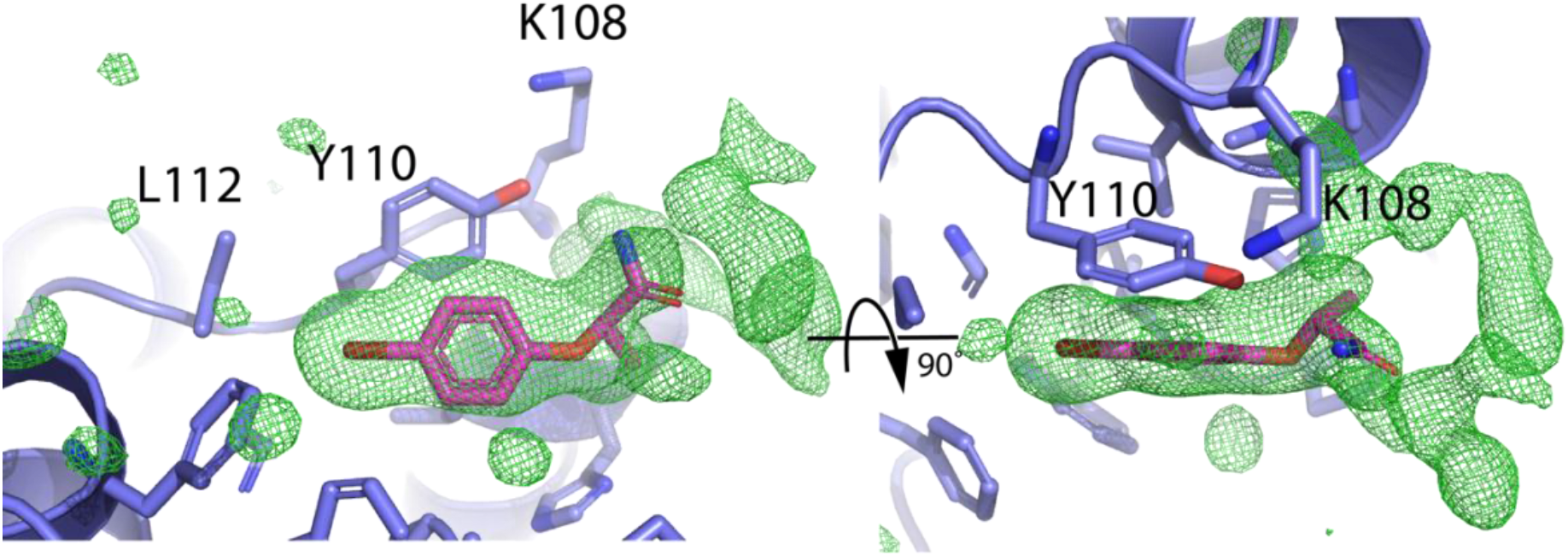
Polder Map around bromophenoxy propanamide (1). Left panel, side view of **1** and corresponding Polder map showing positive Fourier density peak at 3σ surrounding the fragment. Right panel, 90° rotation of the same region. The density on the right of the ligand could be explained by binding of water molecules or of crystallant molecules or a combination thereof.

Binding of fragment **1** caused the side chain of Y110 to shift more than 2 Å from its orientation in the *apo*-structure (measured from the centers of the aromatic rings of the two Y110 conformations), revealing a small hydrophobic pocket (Figure 7). The interactions between **1** and the protein are mostly hydrophobic involving: Y110, W40, F77 and L112. The oxygen of the hydroxy group of Y110 and the nitrogen of **1** are found 3.5 Å apart (Figure 7). Additionally, there are π-stacking interactions between the aromatic rings of the fragment and of Y110 (4.3 Å, measured from the centroid of each ring). The fragment binds within 10 Å of residue C43 (Figure 7) which is part of the redox active site of the protein. During refinement of the structure, the optimal occupancy for the fragment was found to be 0.7, suggesting that the observed density reflects a mixture of the *apo-* and fragment-bound forms of the protein.

**Figure 7 –.**
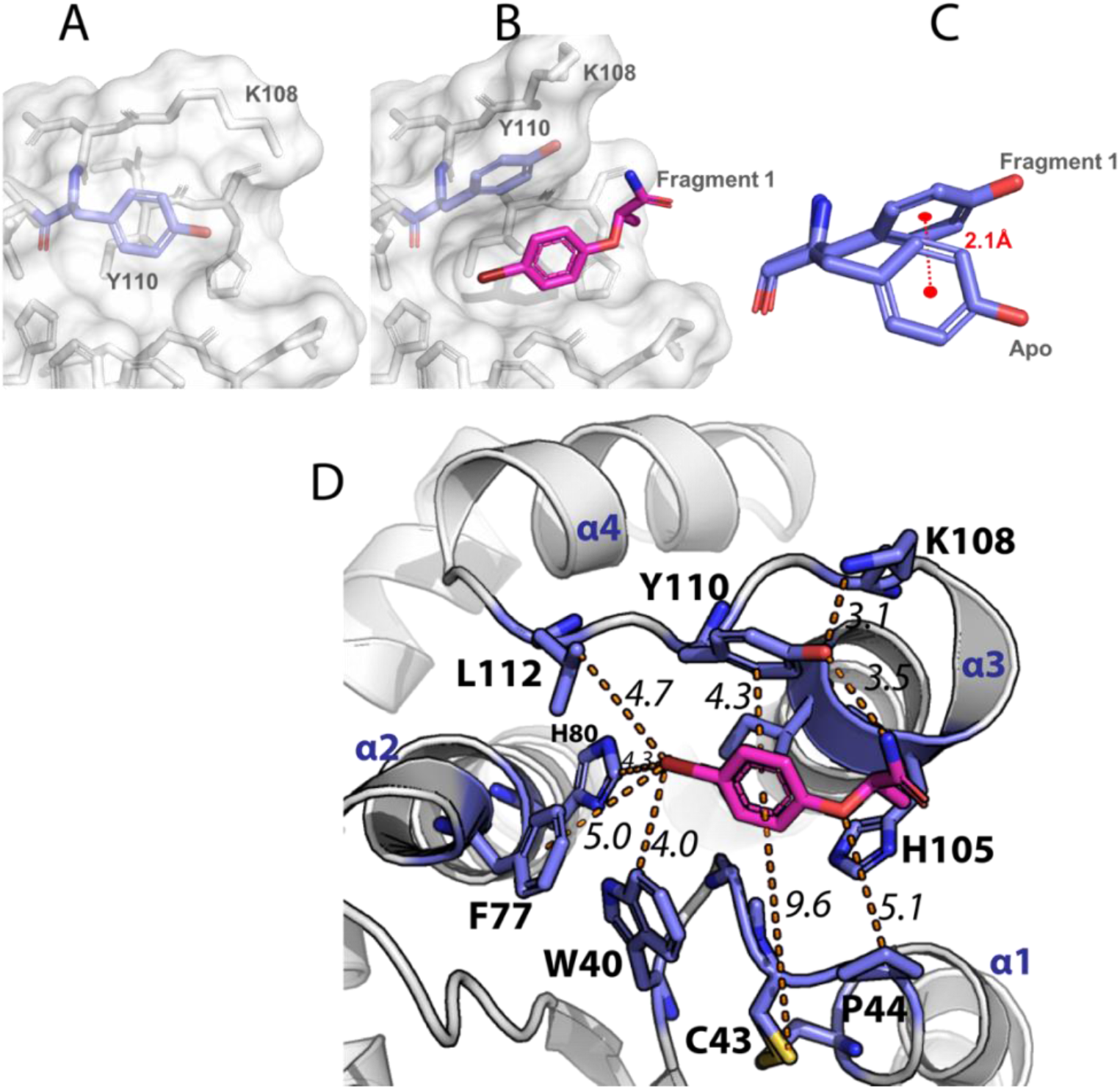
Bromophenoxy propanamide (1) binding requires a shift in Y110 to open a cryptic hydrophobic pocket. **(A)** Side chain position of Y110 in the absence of ligand. **(B)** The same region in presence of **1**, Y110 shifts up (towards the helix α3), opening a binding site for **1**. **(C)** Superposition of Y110 in *apo*- and liganded conformations (Fragment **1**), the center of the benzene ring of the tyrosine is displaced 2.1 Å between the two conformations. **(D)** Structure of BpsDsbA – fragment **1** complex (7LUH) showing the cryptic pocket above the catalytic active site. Fragment **1** (magenta) binds in a hydrophobic pocket. All residues within 5 Å of **1** are shown as purple sticks and labelled. The distances between different atoms or ring centromeres separated by an orange dashed line is given in italics (Angstrom). The sulfurs in the C_43_PHC_46_ active site of BpsDsbA are shown as yellow sticks, the α-helices are also numbered.

### 3.4 BpsDsbA crystallizes as a tetramer when complexed with phenylbenzenesulfonamide (2)

Oxidized BpsDsbA was co-crystallized with **2** in a crystallization solution that typically generated crystals of the *apo*-protein, that is 200 mM Li_2_SO_4_, 100 mM HEPES pH 7.5, 29.5% of PEG 3350. A crystal was harvested and diffracted to a resolution of 2.3 Å in the P 2_1_ space group with unit cell A = 69.4 Å, B = 59.4 Å, C = 105.4 Å and angles α = γ = 90° and β = 104.9° (Table 1). The structure was solved by molecular replacement using the oxidized *apo*-BpsDsbA structure (PDB ID 4K2D, space group P 2_1_2_1_2_1_) as a search model. One solution was found that included four copies of the BpsDsbA per asymmetric unit (Figure 8) and was refined to R_work_ and R_free_ values of 21.27% and 27.04% respectively (Table 1). All four chains of the model align with each other with RMSD between the residues of the different chains below 0.3 Å (alignment of 188 residues with 188 residues for each pairwise comparison), (RMSD chain A - chain B = 0.16 Å, RMSD chain A - chain C = 0.26 Å and RMSD chain A - chain D = 0.25 Å, measured using the Pymol super function (Schödinger, 2015)). The backbone of chain A of this structure also aligns with the *apo*-model PDB ID 4K2D, with a RMSD of 0.24 Å (188 vs 191 residues aligned). The major difference between chain A and the *apo*-model is the truncation of the three N-terminal residues of chain A in the dataset of the complex, relative to the published *apo*-structure, which could not be modelled due to a lack of electron density to justify their placement. In chain D, electron density was poorly resolved for the side chains of residues in the loop 29 to 32 (Figure 8B), and residue Y110 that is reorientated in the presence of **2** (Figure 8C). Modelling of this residue is therefore tentative and must be interpreted with caution.

**Figure 8 –.**
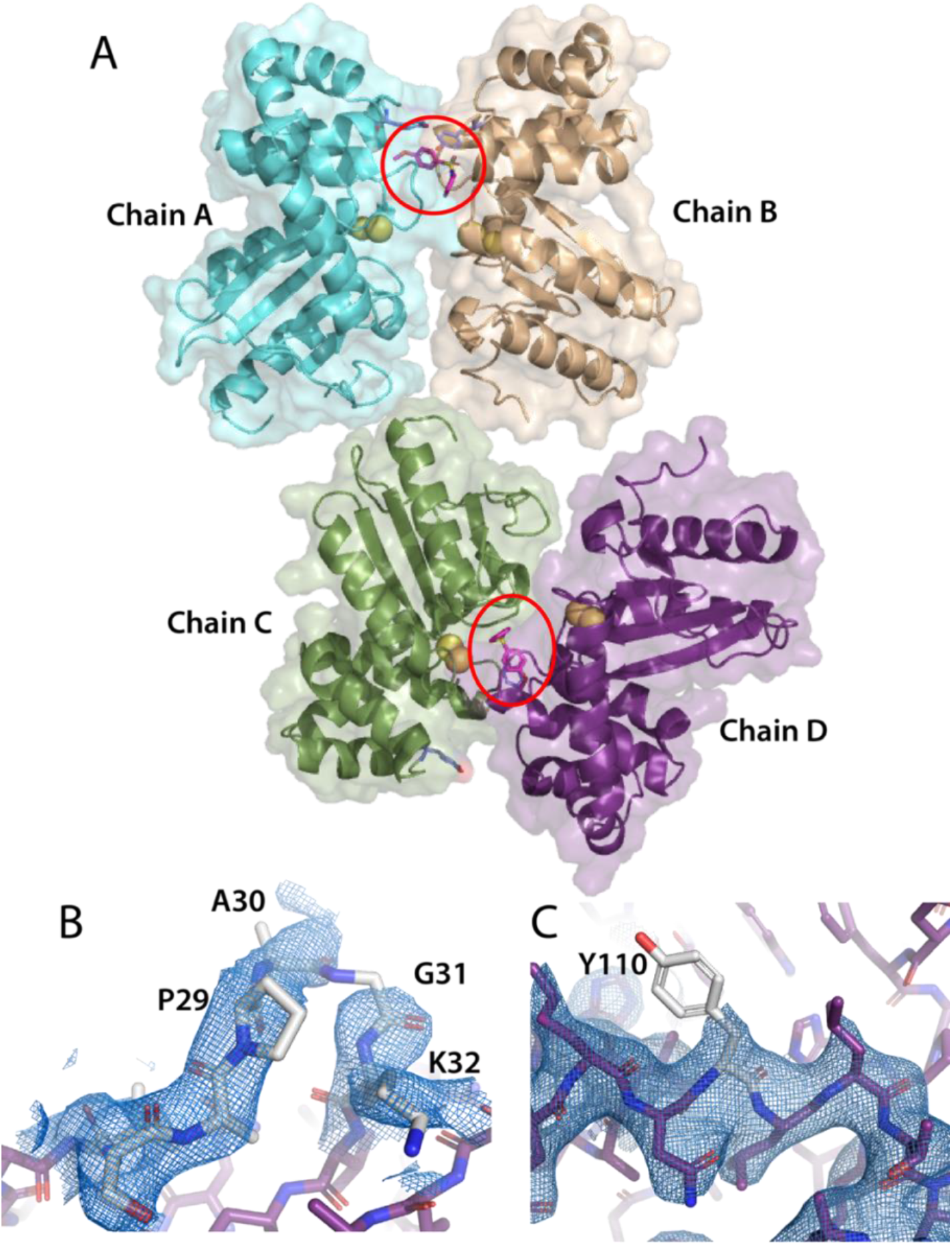
Structure of BpsDsbA crystallized with 2 (PDB ID 7LUJ). **(A)** Overall representation of the four molecules of BpsDsbA in their asymmetric unit. There are two copies of fragment **2**, one binding between chains A and B and the other binding between chains C and D (highlighted by red circles). The active site sulfur atoms are represented as spheres. **(B)** and **(C)** 2Fo-Fc electron densities (at 0.8 σ, in blue) around chain D. **(B)** The map around the loop between residue P29 and K32 (highlighted in grey) was not particularly sharp; single residues were difficult to fit in the densities and the electron density is discontinuous between A30 and G31. **(C)** Similarly, electron density was absent for the side chain of Y110 (note that fragment **2** was removed from this image, for clarity).

Although there are four copies of BpsDsbA in the asymmetric unit, the electron density indicates that only two copies of fragment **2** are bound between the four copies of the protein. One copy of fragment **2** is bound at the interface of chains A and B, its methoxy group binds near Y110 of chain A and its sulfoxide group binds near Y110 of chain B (Figure 9). The second copy of fragment **2** is located at the interface of chains C and D. Again the methoxy side of this copy of the fragment binds near Y110 of chain D while its phenyl side binds closer to the active site on chain C (Figure 9). We note that different parts of this asymmetric fragment are able to bind the same hydrophobic pocket under Y110 of chain A, B and D; this is consistent with the lack of well-defined electrostatic interactions between the fragment and charged residues of the protein. Binding of **2** is apparent in both locations in 2Fo-Fc maps at 1σ (Figure 10A and B), although the density is clearer for the binding site involving chains A and B. The presence of the ligand was confirmed by Polder map analysis which also suggested two alternate bound conformations of **2** at the interface between chains C and D (Figure 10C).

**Figure 9 –.**
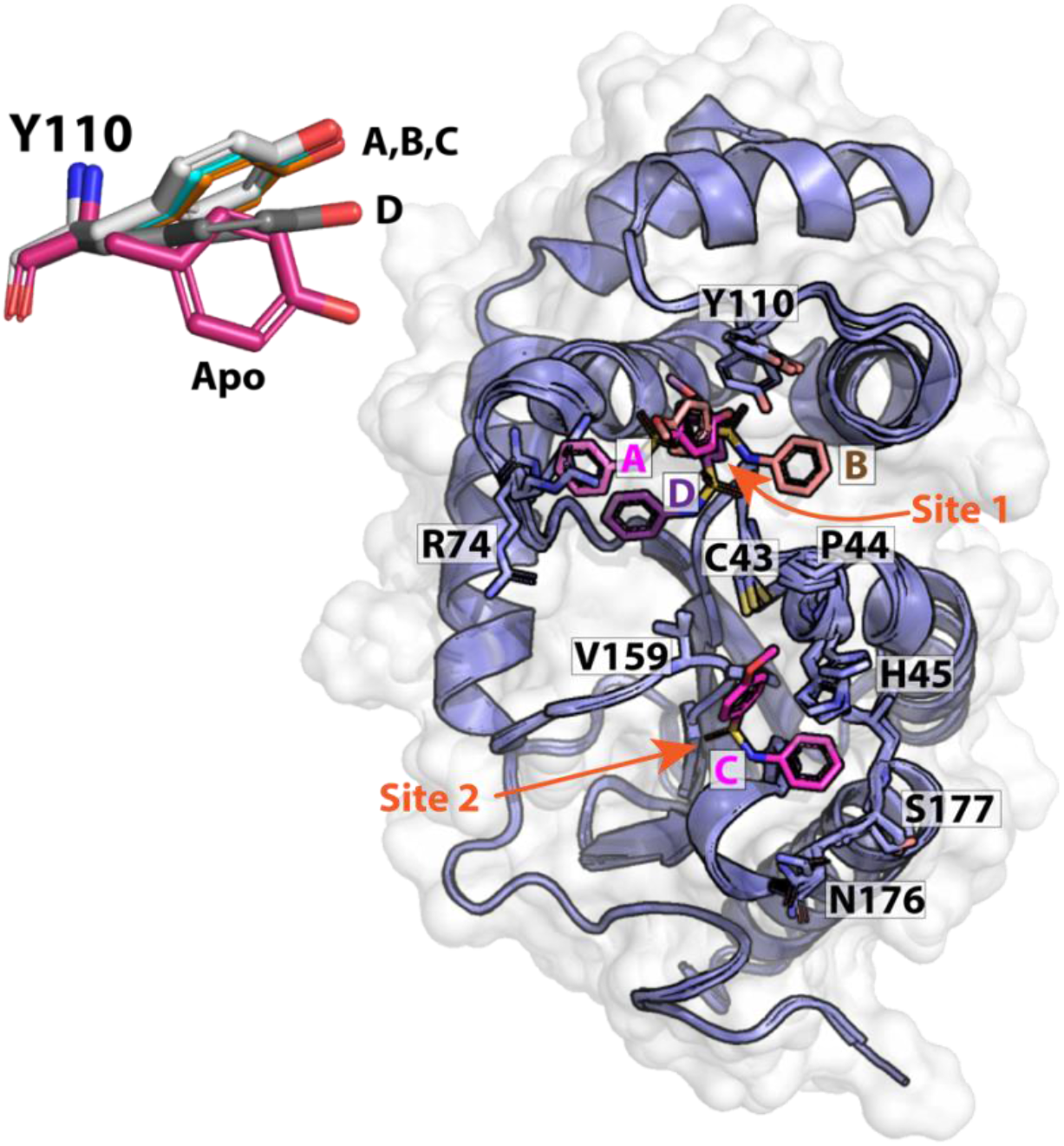
Interaction of phenylbenzenesulfonamide (2) with the four different chains of 7LUJ. In chain A, B and D, **2** binds below Y110 (Site 1, orange arrow). In chain C, the fragment is found closer to the active site (Site 2, orange arrow). Note that there are only two fragments in the asymmetric unit which adopt four different orientations relative to each of the four protein chains (labelled based on the chain they are related to: A, B, C and D). R74 from chain A was also found in a different conformation than in the other chains and wild type protein, making room for **2** to bind to the pocket; other relevant residues are also labelled. The different chains of BpsDsbA are shown as purple cartoon and white surface. The inset figure top left shows the conformations of the four Y110 side chains relative to Y110 of the *apo*-structure. There is more than a 4 Å shift between the hydroxy groups of *apo*-Y110 and Y110 of chain A, B and C.

**Figure 10 –.**
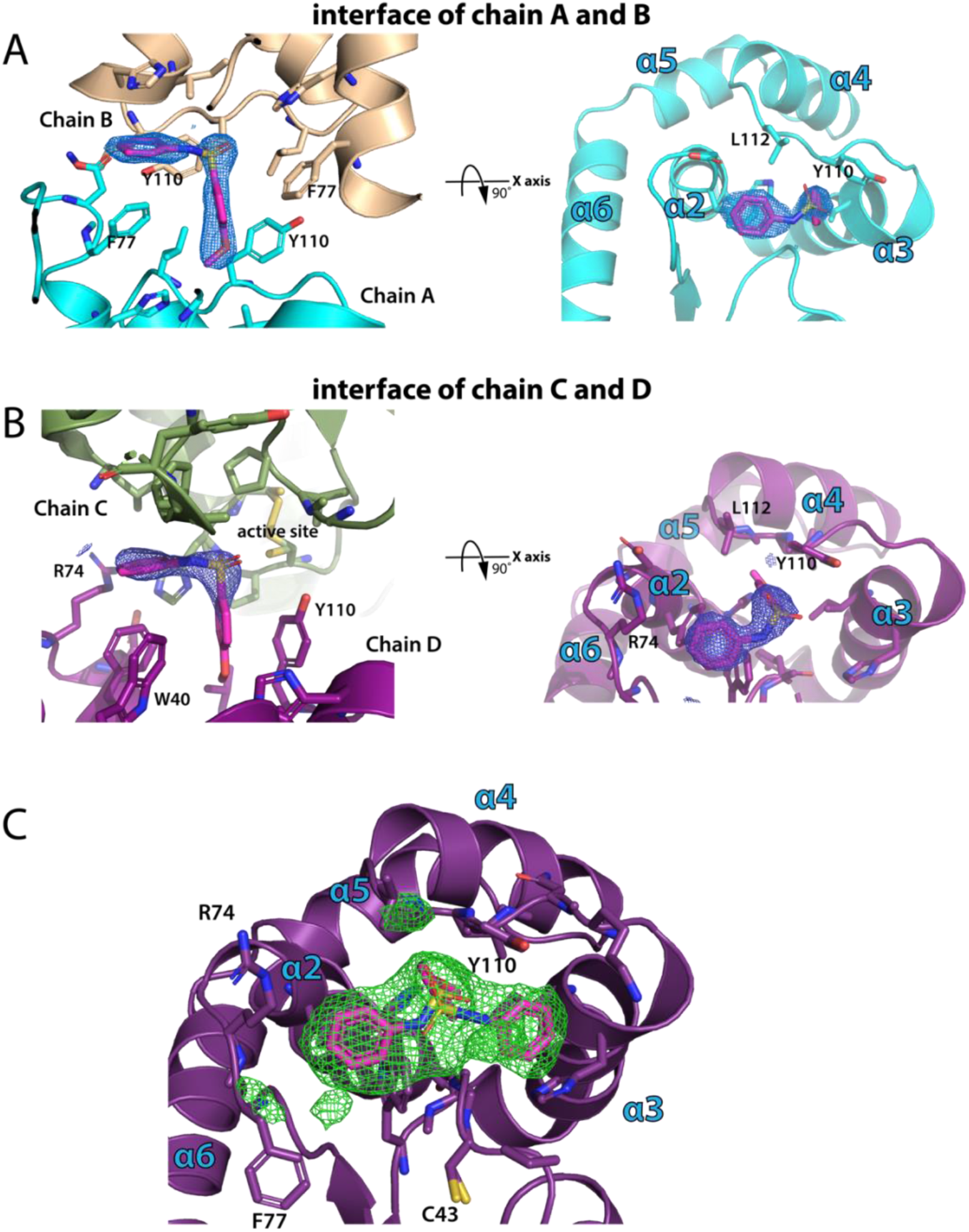
Electron density maps for fragment 2. **(A)** Fragment **2** at the interface of chain A (in cyan) and B (in light brown) is clearly visible on 2Fo-Fc maps set at 1σ in blue suggesting its presence in the pocket. **(B)** 2Fo-Fc maps set at 1σ do not cover the entirety of fragment **2** at the interface of chain C (in olive green) and D (in purple); the methoxy group is not well resolved and its position needed to be confirmed by Polder maps. **(C)** Polder map displayed at 3σ in green, revealed more detail in the density for **2** at the interface of chain C and D, also suggesting that the ligand may bind in two alternate conformations. Residues and helices are labeled where relevant; some residues have been removed from the images for clarity.

It is interesting to note that **2** binds two different sites on the surface of BpsDsbA (Figure 9). On chain A, B and D, fragment **2** is found to bind to the protein in a similar manner to fragment **1**, in a pocket created by the displacement of Y110 towards helix α3, hereafter named site 1 (although the position of Y110 in chain D is not strongly supported by electron densities). In the case of chain C, **2** binds much closer to the active site cysteines, in proximity to residues C43, P44, H45, V159, P160, P172, N176, S177 and L178, hereafter named site 2 (Figure 9). Binding of fragment **2** on site 1 causes a conformational change of chain A’s R74 side chain relative to *apo*-BpsDsbA (Figure 9). Surprisingly Y110 adopts the same orientation in all four molecules in the asymmetric unit whether the fragment is bound or not (fragment **2** does not bind near Y110 of chain C), suggesting that the pocket may not require a ligand to open (Figure 9, inset).

### 3.5 Fragments 1 and 2 do not inhibit the BpsDsbA-BpsDsbB redox cycle

Although the fragments bind weakly to BpsDsbA, we tested whether **1** or **2** were capable of inhibiting the enzymatic activity of BpsDsbA. This was evaluated in a peptide oxidation assay using oxidized BpsDsbA. The assay utilizes a fluorescently-labelled synthetic peptide with cysteines at either end. Oxidation of the substrate by BpsDsbA causes an increase in the fluorescence signal (Halili *et al*., 2015). The reaction was monitored by measuring the increase in fluorescence over the first 10 minutes of the reaction, defined as the initial velocity. Inhibition is indicated by a decrease in the initial velocity compared to the control with no ligand present (addition of matched concentration of DMSO only, Figure S8). Neither of the fragments exhibited any inhibitory activity in this assay; even at a maximum concentration of 20 mM the initial velocity of the reaction was comparable to that of the control reaction. This suggests that the weak binding affinity of the two fragments is not sufficient to compete with or inhibit the peptide used in this assay for binding to BpsDsbA.

## 4. Discussion

DsbA enzymes contribute to the virulence of many Gram-negative bacteria (Coulthurst *et al*., 2008, Heras *et al*., 2009, Ireland *et al*., 2014, McMahon *et al*., 2014) including the often-neglected pathogen *B. pseudomallei*. DsbA proteins have thus been identified as targets for therapeutic drugs (Bocian-Ostrzycka *et al*., 2017, Allen *et al*., 2014, Heras *et al*., 2015, Smith *et al*., 2016).

Several molecules have been reported that inhibit the activity of DsbA enzymes from *E. coli* (Adams *et al*., 2015, Duprez *et al*., 2015, Halili *et al*., 2015), *Pseudomonas aeruginosa* (Mohanty *et al*., 2017) and *Salmonella enterica* serovar Typhimurium (Totsika *et al*., 2018). To date, only one small molecule has been reported to bind to oxidized BpsDsbA and inhibit the enzymatic activity *in vitro* (Nebl *et al*., 2020). BpsDsbA has a shallow hydrophobic groove in comparison to EcDsbA, and a generally flatter surface (McMahon *et al*., 2014), making it a more challenging drug target.

Here we reported the structure and binding interactions of two fragment molecules to oxidized BpsDsbA, both interacting with a small, cryptic pocket close to the protein redox active site. Both fragments, bromophenoxy propanamide (**1)** and 4-methoxy-*N*-phenylbenzenesulfonamide (**2)**, bound under Y110 which was shifted towards helix α3 compared to the *apo*-structure of the protein (Figure 7 and Figure 9). Results were generated using both NMR and X-ray crystallography, and support the findings of Nebl *et al*., (Nebl *et al*., 2020) who had previously identified the presence of a cryptic pocket in the vicinity of W40, C43, C46, R74, I104, Y110 and L112.

Crystallography experiments identified two distinct binding sites on the surface of the protein for **2**. Site 1 is located near Y110 and is replicated in three of the four chains of the ASU of the structure 7LUJ (chain A, B and D). This site is also supported by the NMR experiments which show a large CSP for Y110 upon addition of **2** to the protein; and by the initial crystal soaks analyzed with PanDDA (Figure 1). The second site (site 2) is only visible on chain C of 7LUJ and NMR experiments do not show clear CSP for residues adjacent to site 2 suggesting that this site might be an artefact of crystallization.

The binding of fragments **1** and **2** to BpsDsbA is weak (K_D_ > 2 mM) consistent with their small size. Weak binding is evident in the partial occupancy and high B-factors of the modelled ligand in the crystal structures (Table 1). We found very weak or no evidence of fragment binding to reduced BpsDsbA (Figure S7), supporting the hypothesis that the redox state of BpsDsbA governs the formation of the cryptic pocket, and perhaps the flexibility of the Y110 side chain, which allows the pocket to form (Nebl *et al*., 2020).

The identification and characterization of the binding of fragments **1** and **2** to BpsDsbA is a key first step towards understanding this cryptic pocket and the dynamic behavior of the active site at atomic resolution. This pocket is of interest because of its proximity to the active site, which suggest that expanding these fragments may generate more potent compounds that block the active site, and inhibit the activity of BpsDsbA. The results presented here provide a starting point for the elaboration and further optimization of more potent small molecule inhibitors for BpsDsbA using rational drug design.

## 5. Acknowledgments

The authors acknowledge the University of Queensland Remote Operation Crystallization and X-ray diffraction facility (UQ ROCX) part of the Microscopy Australia Facility at the Centre for Microscopy and Microanalysis.

The authors also acknowledge use of the Australian Synchrotron (ANSTO), in particular the MX2 beamline (McPhillips *et al*., 2002) and its detector which is supported by the Australian Cancer Research Foundation.

The National Deuteration Facility is partly funded by the National Collaborative Research Infrastructure Strategy (NCRIS), an Australian Government initiative.

We gratefully acknowledge the support of the Griffith University eResearch Services Team and the use of the High Performance Computing Cluster “Gowonda” to complete this research

