## Supporting Info S1 - S8 for "Identification and characterization of two drug-like fragments that bind to the same cryptic binding pocket of *Burkholderia pseudomallei* DsbA"

### Supporting information

#### Synthesis of 4-Methoxy-*N*-phenylbenzenesulfonamide

General experimental detail:

All reactions were carried out in dry solvents under anhydrous conditions unless otherwise stated. All chemicals were purchased from commercial suppliers and used without further purification. All reactions were monitored by TLC using silica plates with visualization of eluted bands by UC fluorescence ($\lambda$ = 254 nm) and charring with vanillin stain (6 g vanillin in 100 mL of EtOH containing 1% v/v 98% Sulfuric acid. Silica gel flash chromatography was performed using silica gel 60 Å (230-400 mesh). NMR (1D $^{1}$H, 1D $^{13}$C, 2D [$^{1}$H,$^{1}$H] - COSY, 2D [$^{13}$C, $^{1}$H] - HSQC and 2D [$^{13}$C, $^{1}$H] - HMBC) spectra were recorded on a Bruker AVANCE III HD 500 MHz NMR spectrometer equipped with a BBO probe at 25 ˚C. Chemical Shifts for $^{1}$H and $^{13}$C NMR obtained in Chemical Shifts for $^{1}$H and $^{13}$C NMR obtained in CDCl3 are reported in ppm relative to residual solvent proton ($\delta$ = 7.26 ppm) and carbon ($\delta$ = 77.16 ppm) signals, respectively. Signal splitting multiplicity is indicated as follows: s (singlet), d (doublet), t (triplet), q (quartet), m (multiplet), dd (doublet of doublets), and br (broad signal).

**4-Methoxy-*N*-phenylbenzenesulfonamide** (**2**): The titled fragment was prepared according to literature procedures (Bernar *et al.*, 2018) from 4-methoxybenzenesulfonyl chloride (0.250 g, 1.21 mmol, 1 equiv.) and aniline (0.120 mL, 1.33 mmol, 1.1 equiv.) in a mixture of pyridine (0.195 mL, 2.42 mmol, 2 equiv.) and $CH_2Cl_2$ (2.85 mL). The crude product was purified by silica gel flash chromatography employing a mobile phase gradient of 20% to 40% EtOAc in *n*-hexane to afford the desired product as a slightly yellow viscous oil (0.2967 g, 93% yield) which solidified to a white solid upon storage at -20 °C. $^{1}$H NMR and $^{13}$C NMR spectroscopic data were consistent with reported values.(Bernar *et al.*, 2018) $^{1}$H NMR (500 MHz, $CDCl_3$) $\delta_H$ 7.76 – 7.70 (m, 2H), 7.25 – 7.20 (m, 2H), 7.13 – 7.06 (m, 3H), 7.02 (br s, 1H), 6.91 – 6.86 (m, 2H), 3.81 (s, 3H). $^{13}$C NMR (125 MHz, $CDCl_3$) $\delta_C$ 163.2, 136.8, 130.7, 129.6, 129.4, 125.4, 121.6, 114.3, 55.7.

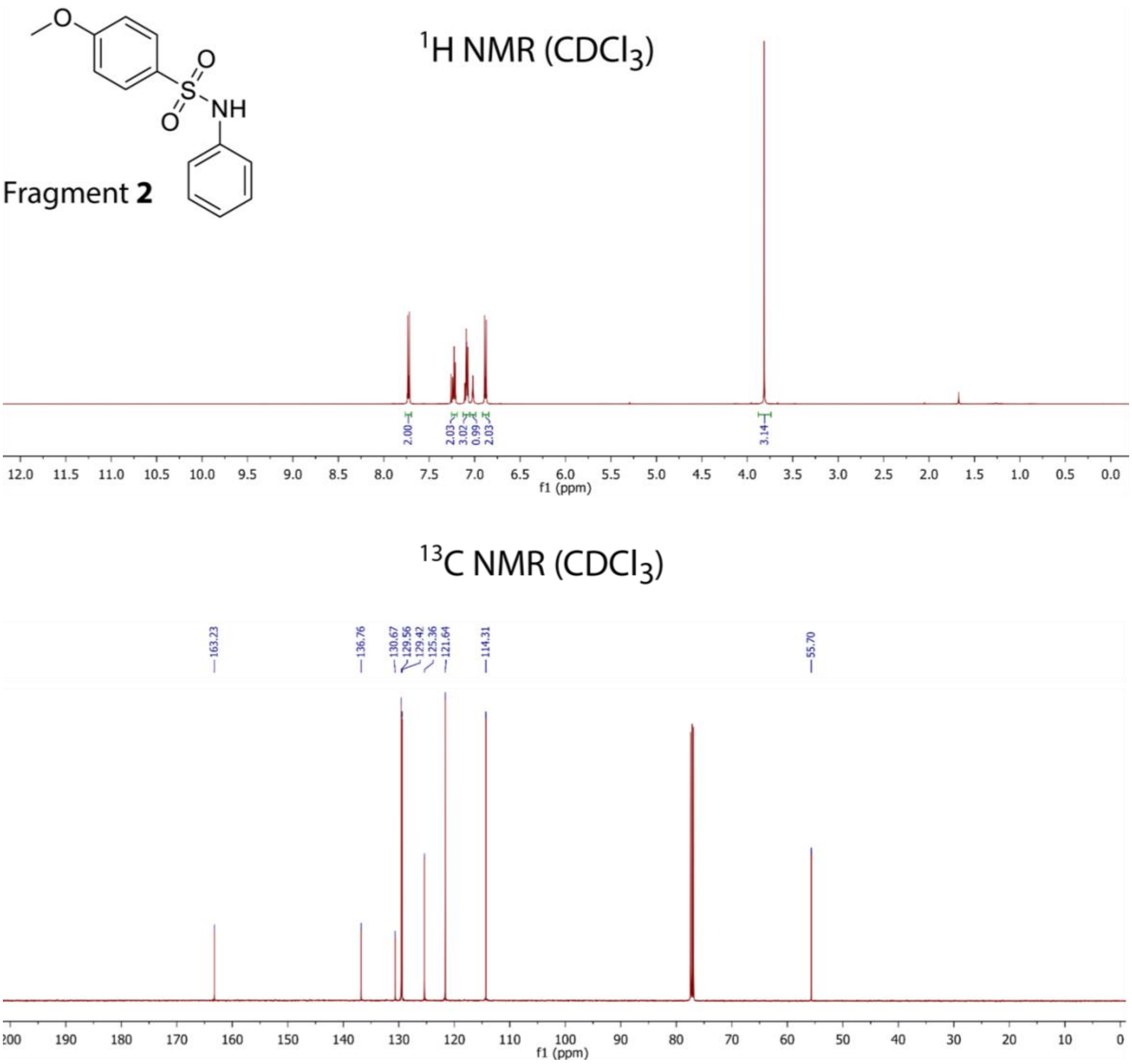


**S1 – NMR spectra of fragment 2.** Spectra acquired at 500 MHz for hydrogen NMR and 125 MHz for Carbon NMR

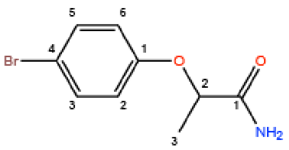

2-(4-bromophenoxy)propanamide (**1**)

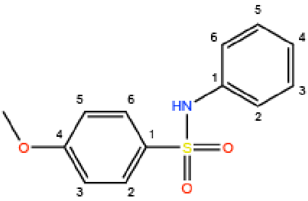

4-methoxy-*N* -phenylbenzenesulfonamide (**2**)

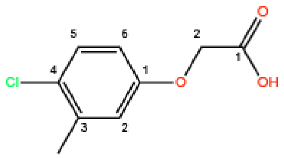

2-(4-chloro-3-methylphenoxy)acetic acid

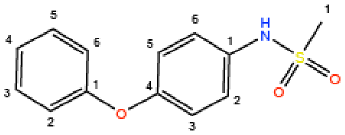

*N*-(4-phenoxyphenyl)methanesulfonamide

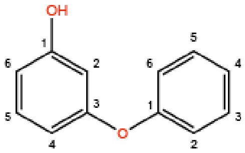

3-phenoxyphenol

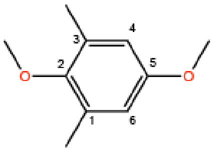

2,5-dimethoxy-1,3-dimethylbenzene

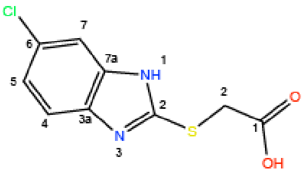

2-((6-chloro-1*H*-benzo[*d*]imidazol-2-yl)thio)acetic acid

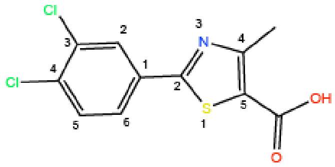

2-(3,4-dichlorophenyl)-4-methylthiazole-5-carboxylic acid

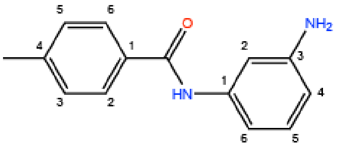

*N* -(3-aminophenyl)-4-methylbenzamide

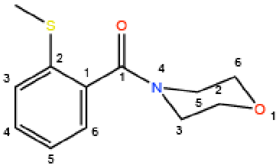

(2-(methylthio)phenyl)(morpholino)methanone

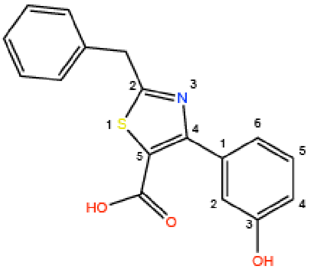

2-benzyl-4-(3-hydroxyphenyl)thiazole-5-carboxylic acid

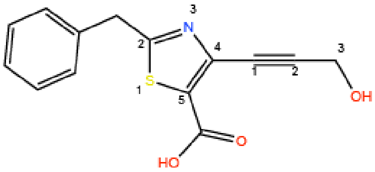

2-benzyl-4-(3-hydroxyprop-1-yn-1-yl)thiazole-5-carboxylic acid

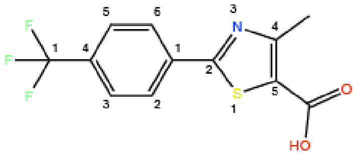

4-methyl-2-(4-(trifluoromethyl)phenyl)thiazole-5-carboxylic acid

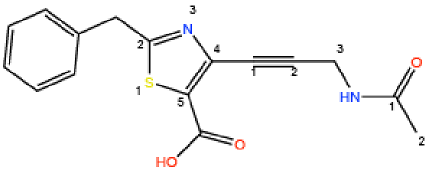

4-(3-acetamidoprop-1-yn-1-yl)-2-benzylthiazole-5-carboxylic acid

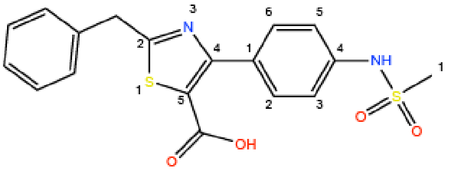

2-benzyl-4-(4-(methylsulfonamido)phenyl)thiazole-5-carboxylic acid

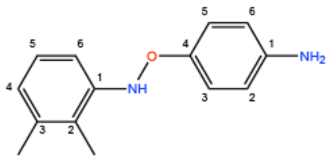

4-(((2,3-dimethylphenyl)amino)oxy)aniline

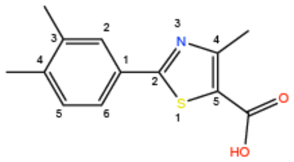

2-(3,4-dimethylphenyl)-4-methylthiazole-5-carboxylic acid

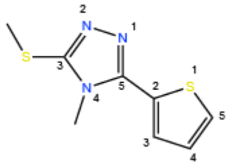

4-methyl-3-(methylthio)-5-(thiophen-2-yl)-4*H*-1,2,4-triazole

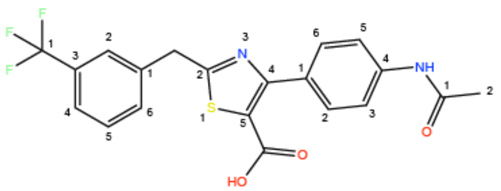

4-(4-acetamidophenyl)-2-(3-(trifluoromethyl)benzyl)thiazole-5-carboxylic acid

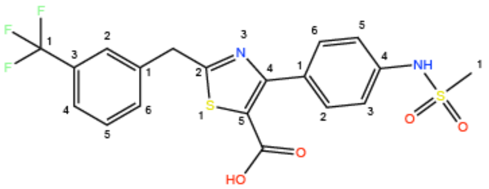

4-(4-(methylsulfonamido)phenyl)-2-(3-(trifluoromethyl)benzyl)thiazole-5-carboxylic acid

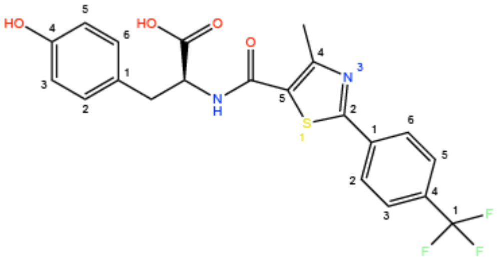

(4-methyl-2-(4-(trifluoromethyl)phenyl)thiazole-5-carbonyl)-*L*-tyrosine

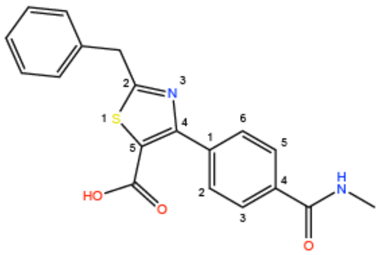

2-benzyl-4-(4-(methylcarbamoyl)phenyl)thiazole-5-carboxylic acid

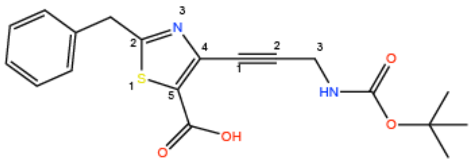

2-benzyl-4-(3-((*tert*-butoxycarbonyl)amino)prop-1-yn-1-yl)thiazole-5-carboxylic acid

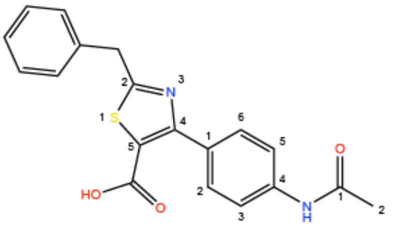

4-(4-acetamidophenyl)-2-benzylthiazole-5-carboxylic acid

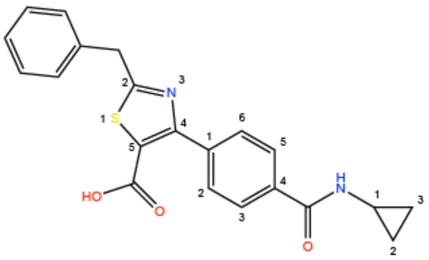

2-benzyl-4-(4-(cyclopropylcarbamoyl)phenyl)thiazole-5-carboxylic acid

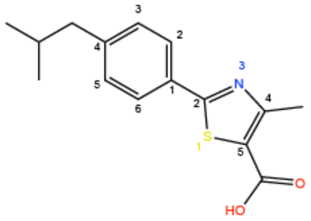

2-(4-isobutylphenyl)-4-methylthiazole-5-carboxylic acid

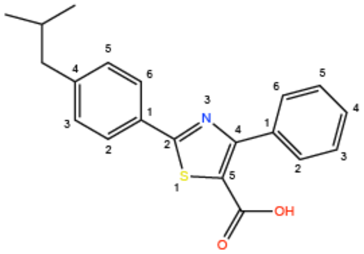

2-(4-isobutylphenyl)-4-phenylthiazole-5-carboxylic acid

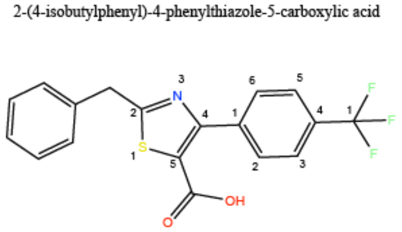

2-benzyl-4-(4-(trifluoromethyl)phenyl)thiazole-5-carboxylic acid

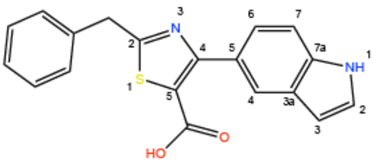

2-benzyl-4-(1*H*-indol-5-yl)thiazole-5-carboxylic acid

**S2: Chemical structure of the 29 unique fragments tested in X-ray crystallization experiments in this study**

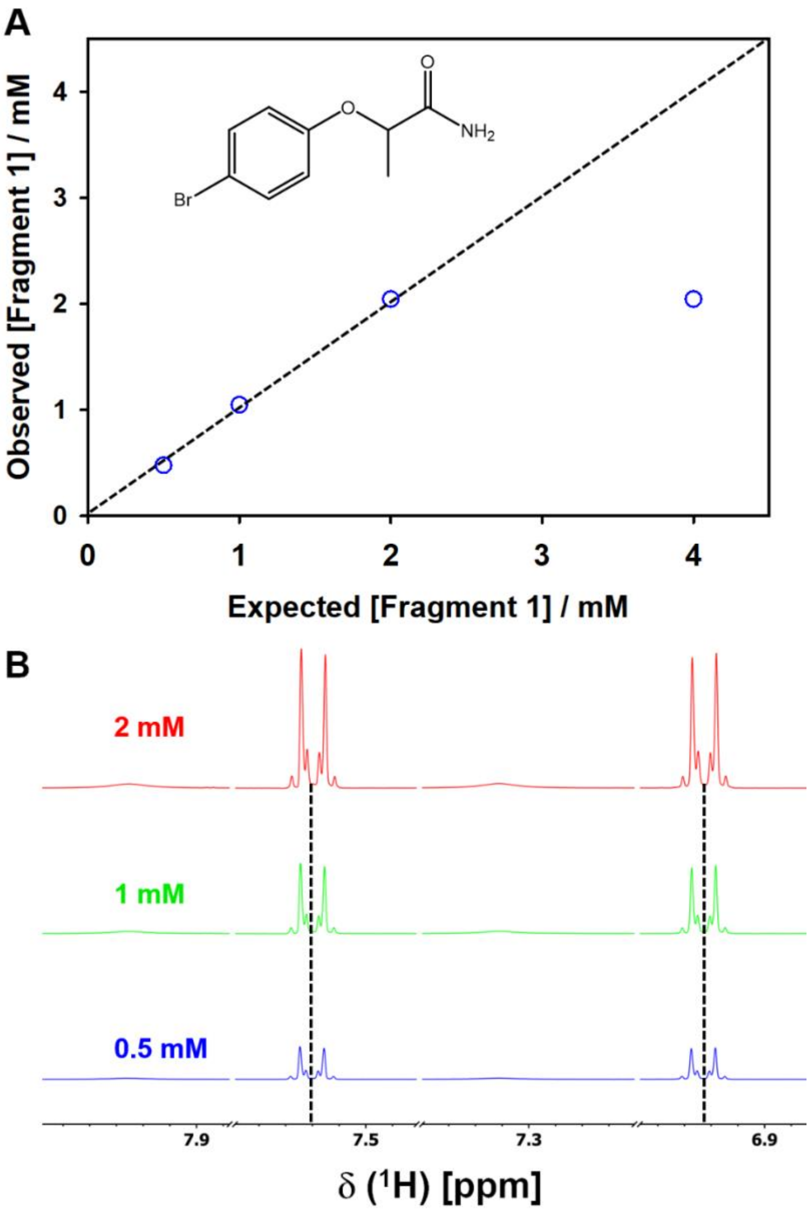


**S3 – Solubility assessment of bromophenoxy propanamide (1) in aqueous NMR buffer**. **(A)** Correlation between calculated and measured concentration of **1**. Dotted diagonal line represents the linear relationship between the expected and observed values. Molecular structure of **1** is inset. **(B)** Zoomed region of the 1D $^{1}$H spectra showing the aromatic resonances of **1** to highlight the increasing peak intensity resulting from increasing concentration. At 4 mM, precipitate was observed. As such a large variance was observed between the expected and observed concentration, the corresponding 1D $^{1}$H spectrum has been excluded in (B). This suggests that **1** is soluble at concentrations $\leq$ 2 mM in aqueous NMR buffer at 298 K.

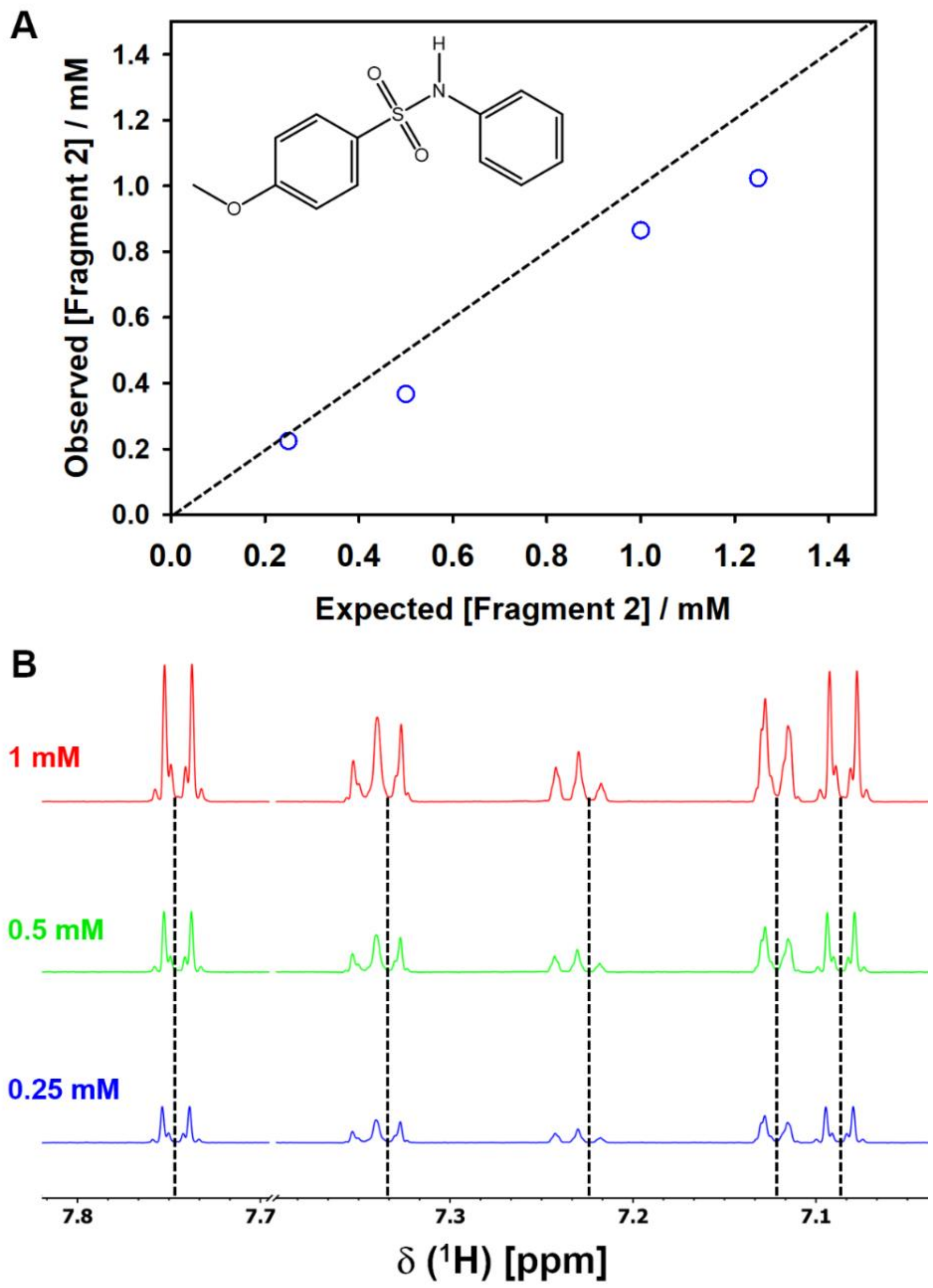


**S4 – Solubility assessment of methoxy-phenylbenzenesulfonamide (2) in aqueous NMR buffer.** (**A**) Correlation between expected and observed concentration of **2.** Molecular structure of **2** is inset. Dotted diagonal line represents the linear relationship between the calculated and measured values. (**B)** Zoomed region of the 1D $^{1}$H spectra showing the aromatic resonances of **2** to highlight the increasing peak intensity resulting from increasing concentration. At 1.25 mM visible precipitate was observed, the corresponding 1D $^{1}$H spectrum has been excluded in (B). These data suggest that **2** is soluble at concentrations ≤ 1 mM in aqueous NMR buffer at 298 K.

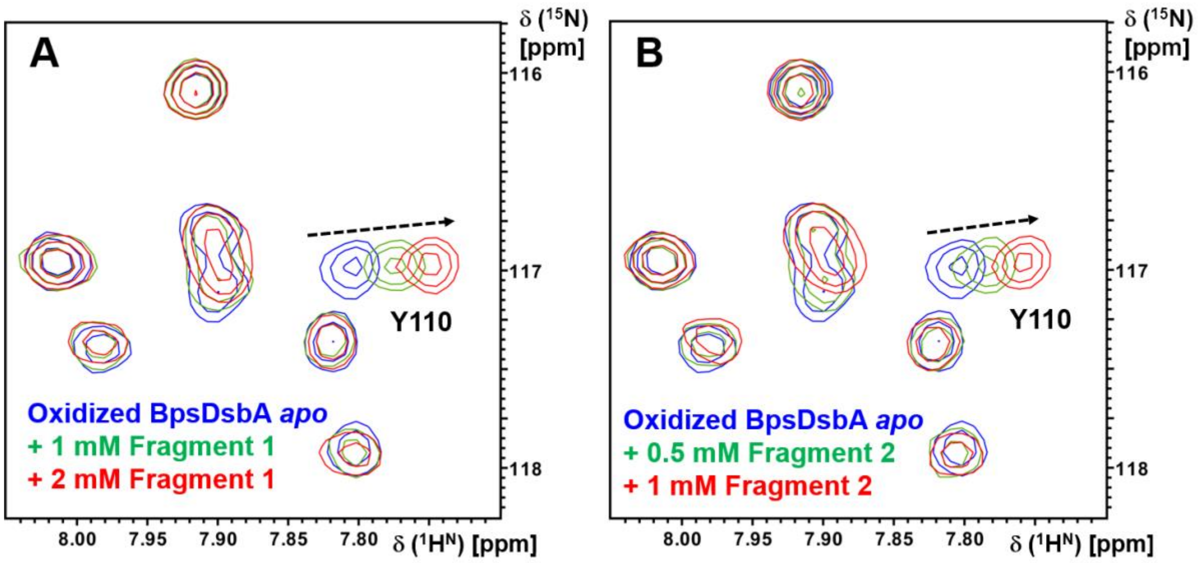


**S5 – Characterization of 1 and 2 binding to oxidized BpsDsbA by 2D [$^{15}$N, $^{1}$H]-HSQC NMR.** (**A**) 2D [$^{15}$N,$^{1}$H]-HSQC overlay of 100 μM [U-$^{15}$N]- labelled oxidized BpsDsbA with increasing concentrations of **1.** **(B)** 2D [$^{15}$N,$^{1}$H]-HSQC overlay of 100 μM [U-$^{15}$N]-labelled oxidized BpsDsbA with increasing concentrations of **2.** Concentration dependent backbone amide chemical shift perturbation of Y110 is highlighted in each case.

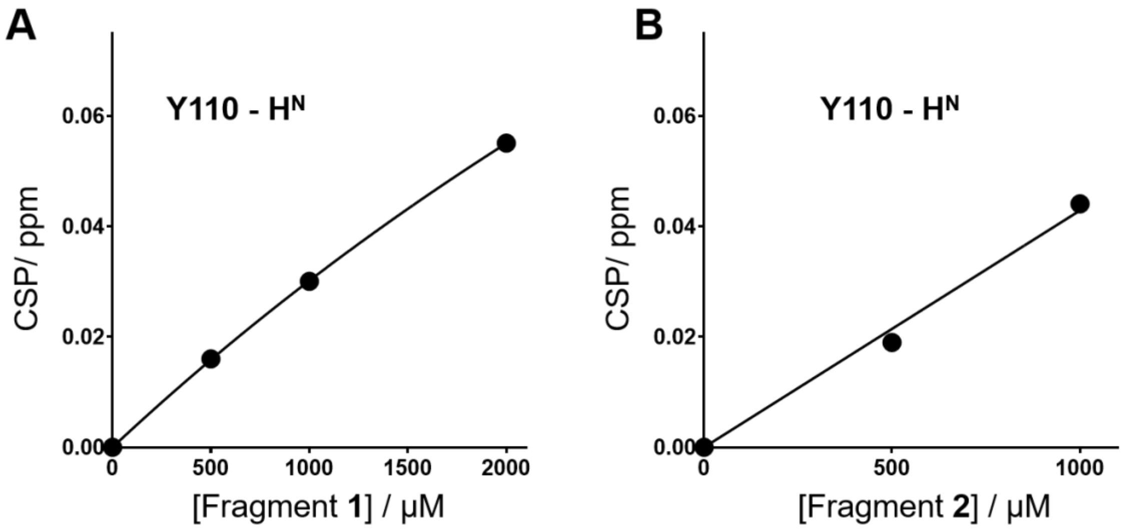


**S6. Characterization of fragment 1 and 2 binding affinities to oxidized BpsDsbA by 2D [$^{15}$N, $^{1}$H]-HSQC NMR.** (**A**) Y110 backbone amide CSP resulting from the addition of 0.5 mM,1 mM and 2 mM fragment **1**. **(B)** Y110 backbone amide CSP resulting from the addition of 0.5 mM and 1 mM fragment **2**. CSP responses were observed to increase linearly with concentration for both fragments and reliable estimates of $K_D$ could not be obtained. These data indicate weak interaction between fragments and oxidized BpsDsbA.

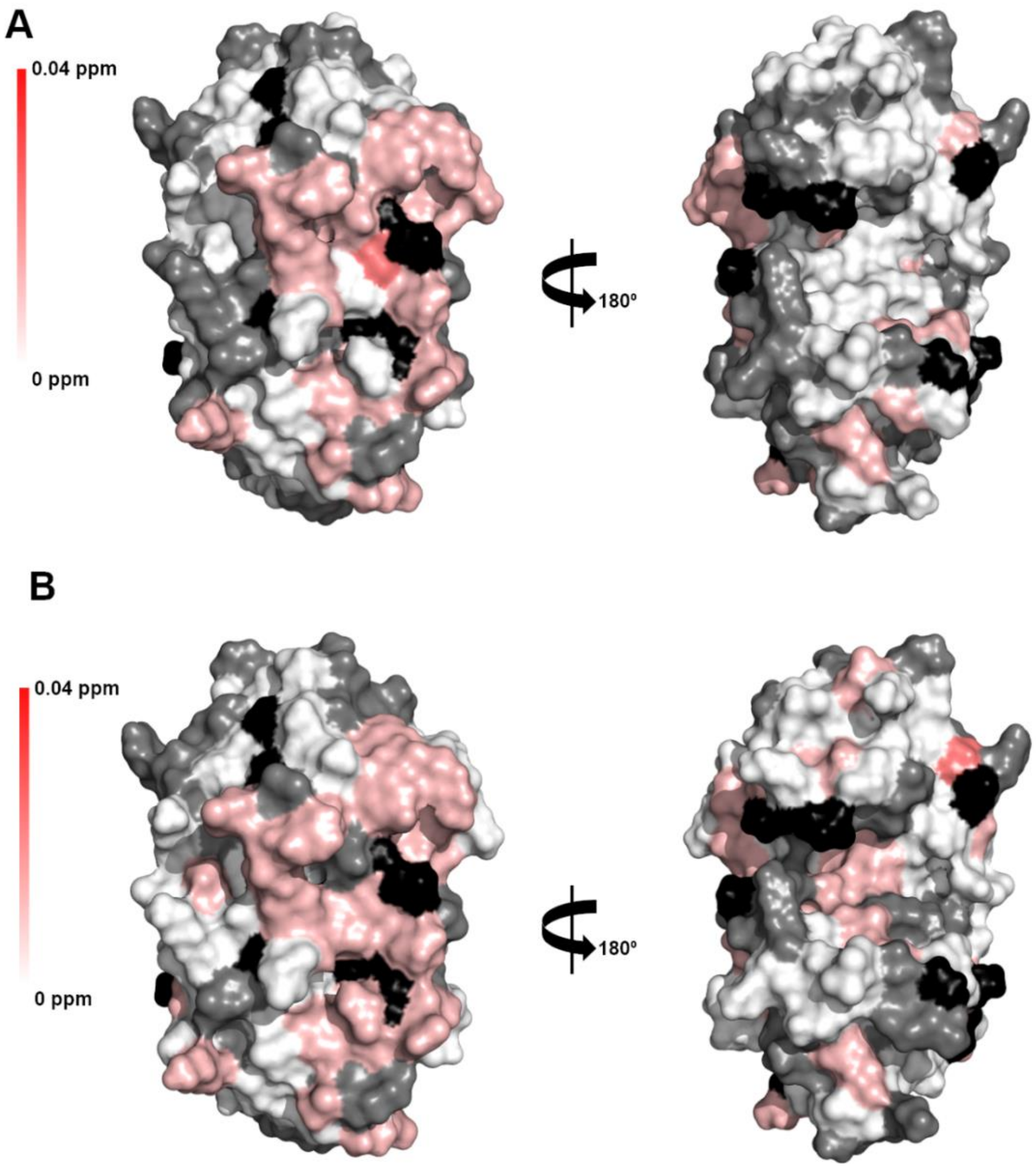


**S7. Characterization of fragment 1 and 2 binding to reduced BpsDsbA by 2D [$^{15}$N, $^{1}$H]-HSQC NMR. (A)** CSPs resulting from the addition of 1 mM of fragment **1** are mapped onto the crystal structure of oxidized BpsDsbA (PDB ID: 4K2D) as a color gradient from red (CSP = 0.04 ppm) to white (CSP = 0 ppm). **(B)** CSPs resulting from the addition of 1 mM of fragment **2** are mapped onto the crystal structure of oxidized BpsDsbA (PDB ID: 4k2d) as a color gradient from red (CSP = 0.04 ppm) to white (CSP = 0 ppm). Residues with unassigned backbone amides and proline residues are shown in black. Non-shifting residues are shown in grey. N-terminal residues were removed for clarity. Three-dimensional structure of reduced BpsDsbA is not solved to date therefore we used the crystal structure of oxidized BpsDsbA for the CSP mapping analysis.

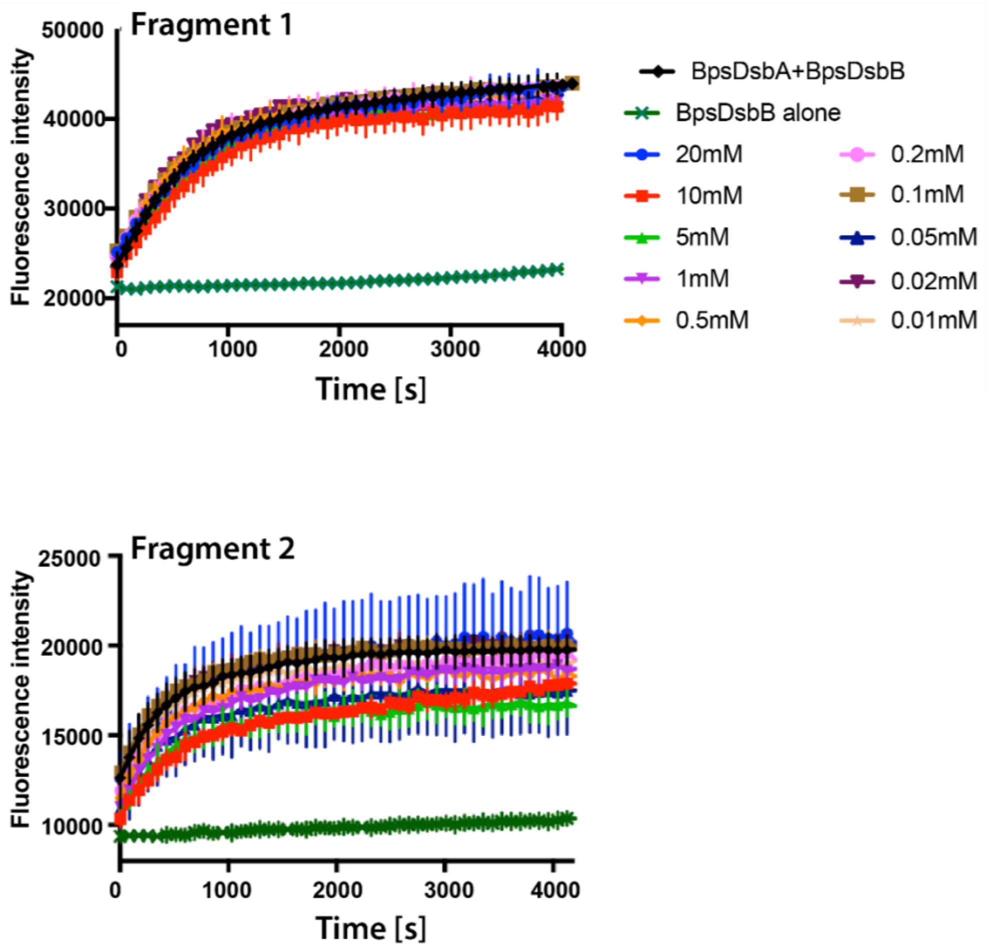


**S8: Peptide oxidation assay in the presence of fragment 1 (upper panel) or 2 (lower panel).** The fragments did not inhibit enzyme activity in this assay, even at a concentration up to 20 mM. The fragments **1** and **2** were dissolved in DSMO and mixed with the protein to the final concentration indicated in the legend. Each different trace corresponds to the fluorescence value of two technical replicates at each concentration, and the error bar depicts the standard error. The dark green trace corresponds to the reaction without BpsDsbA (negative control, representing an entirely inhibited reaction) and the black trace corresponds to an uninhibited reaction (positive control).
